# BarTn7: Optimizing Bacterial Lineage Tracking at Sub-Species Resolution for Population Dynamics in Ecological and Evolutionary Studies

**DOI:** 10.1101/2025.11.24.690285

**Authors:** Robin A. Herbert, Adam M. Deutschbauer

## Abstract

Communities of bacteria undergo population bottlenecks which are crucial to their population, ecological, and evolutionary dynamics. However, conventional amplicon sequencing cannot distinguish such demographic shifts between very closely related lineages. Here we describe BarTn7, an optimized method for bacterial lineage tracking through chromosomal integration of phenotypically neutral DNA barcodes with transposon Tn7. By combining conventional conjugative plasmids into a single vector and leveraging a parts-based strategy to optimize delivery to different recipients, BarTn7 increases barcoding efficiency and enables the systematic application of this tool to diverse bacteria. We tested BarTn7 in multiple bacterial species and confirmed a lack of lineage-specific growth effects. We then used BarTn7 to measure the colonization of *Panicum virgatum* roots under differing phosphate concentrations. Lineage tracking enabled discrimination between unique colonization events and the proliferation of existing bacteria during root colonization, the ratio of which differed between three plant-associated bacterial species. The strategy further allowed the measurement of phosphate-dependent ingress rates of the root endosphere by *Paraburkholderia phytofirmans* PsJN. We then demonstrated the effectiveness of BarTn7 to detect adaptive mutation(s) and facilitate identification of mutant lineages. Additionally, we demonstrated that BarTn7 is more accurate than conventional 16S rRNA gene sequencing at measuring community composition of a 5-member synthetic bacterial community (SynCom) and compares favorably to shotgun metagenomics. Our results illustrate the utility of BarTn7 as a simple, cost-efficient, and broadly applicable method of measuring bacterial population dynamics at sub-species resolution.

## Introduction

Many microbial communities undergo demographic shifts at the strain level which are ecologically relevant and impact host physiology (**1–4**). During colonization of the plant root, for instance, bacteria must first move towards and attach to the root, then penetrate root tissue before proliferating therein (**5–6**). Each of these steps imposes a population bottleneck, however the relative contributions of ingress and proliferation within tissue are difficult to measure. 16S rRNA gene sequencing is generally limited to measuring these shifts at the level of bacterial genera and can be biased towards specific taxa (**7–8**). In contrast, whole-genome metagenomics can measure genetic diversity at high resolution but cannot assign alleles to specific strains. The ability to measure microdiversity quickly and cost-efficiently would therefore enable greater understanding of bacterial population dynamics.

To make these measurements, we describe BarTn7, a scalable method of quantifying lineage dynamics within populations of clonal bacteria. BarTn7 uses the previously described ‘magic-pool’ method of adding DNA barcodes and other genetic cargo in diverse bacteria (**9**) in a Tn7 transposon system. The Tn7 transposon is ideal for high-resolution measurement of nearly isogenic populations of bacteria, as the TnsD subunit of Tn7 protein targets transposition downstream of a highly conserved gene, *glmS* (**10**), such that BarTn7 is applicable to a variety of bacteria. Targeted integration downstream of *glmS* typically does not introduce biased fitness effects between recipient strains. Furthermore, the presence of an initial Tn7 transposon has been shown to prevent subsequent transposition events (**11**), such that the gene cargo can be delivered in single copy.

We introduced a DNA-barcoded Tn7 transposon encoding an antibiotic resistance marker and fluorophore into the genomes of diverse recipient bacteria. Transposition downstream of *glmS* was validated by PCR in all species, including those with multiple homologs of the gene. We used this system to generate populations of thousands to hundreds-of-thousands of barcoded lineages within seven individual bacterial species and passaged these libraries to confirm that lineages grow without bias relative to their broader community. We then used barcoded libraries of three root-associated bacteria to measure the colonization of *Pannicum virgatum* (henceforth “switchgrass”) roots under two phosphorus concentrations. Our measurements demonstrate that root colonization by different species can be driven by increased proliferation or additional recruitment from the surrounding medium. Furthermore, we show that low phosphate causes a change in strain recruitment of these bacteria such that the sampled communities are less evenly distributed–with individual lineages becoming more prolific.

To confirm the utility of BarTn7 in ecological studies, we grew a synthetic community (SynCom) of five barcoded bacterial species in four different media and compared their community composition as measured by BarTn7, 16S rRNA gene amplification, and Illumina shotgun metagenomics. In these experiments, BarTn7 was more accurate than 16S sequencing even when controlling for 16S gene copy number, comparably accurate to WGS, and less costly than both. As a proof of concept, we also used BarTn7 to predict beneficial mutations within barcoded populations of *Escherichia coli* BW25113 and *Klebsiella michiganensis* M5a1 treated with trimethoprim.

Finally, we optimized the Tn7 delivery vector by testing different cargo orientations, combining the transposase enzymes and transposon into a single vector, and implementing a parts-based method to identify genetic components which work best for a given recipient. These alterations served to increase conjugative efficiency, reduce unwanted toxicity of the system to certain recipient hosts, and rapidly identify the ideal combination of genetic parts for BarTn7 implementation for several otherwise recalcitrant bacteria. Using this BarTn7 system we were able to generate large, barcoded communities in multiple bacteria and use these to draw novel inferences around their population, ecological, and evolutionary dynamics.

## Results

### Overview of BarTn7

Our BarTn7 approach is summarized in **Fig. 1**. Incorporation of Tn7 typically uses a two-plasmid system wherein one vector contains the transposon, and another encodes the transposase (**12–14**). Briefly, we constructed a vector pool, pRH05, each of which contains a kanamycin resistance cassette, GFP, and a random 20 nucleotide DNA barcode, flanked by the Tn7 left and right arms (**Fig. 1A**). The resulting plasmid pool contained ∼4 million unique barcodes and was used for the initial tests of the BarTn7 system. We performed a triparental conjugation of this barcoded transposon library and the Tn7 transposase encoded on a separate “helper” plasmid pTNS2 into 7 individual recipient ba-cteria (**Fig. 1B**). For each target bacterium, we validated the expected locus and orientation of transposition by PCR targeting the terminus of *glmS* (**Fig. 1C**).

**Figure 1.**
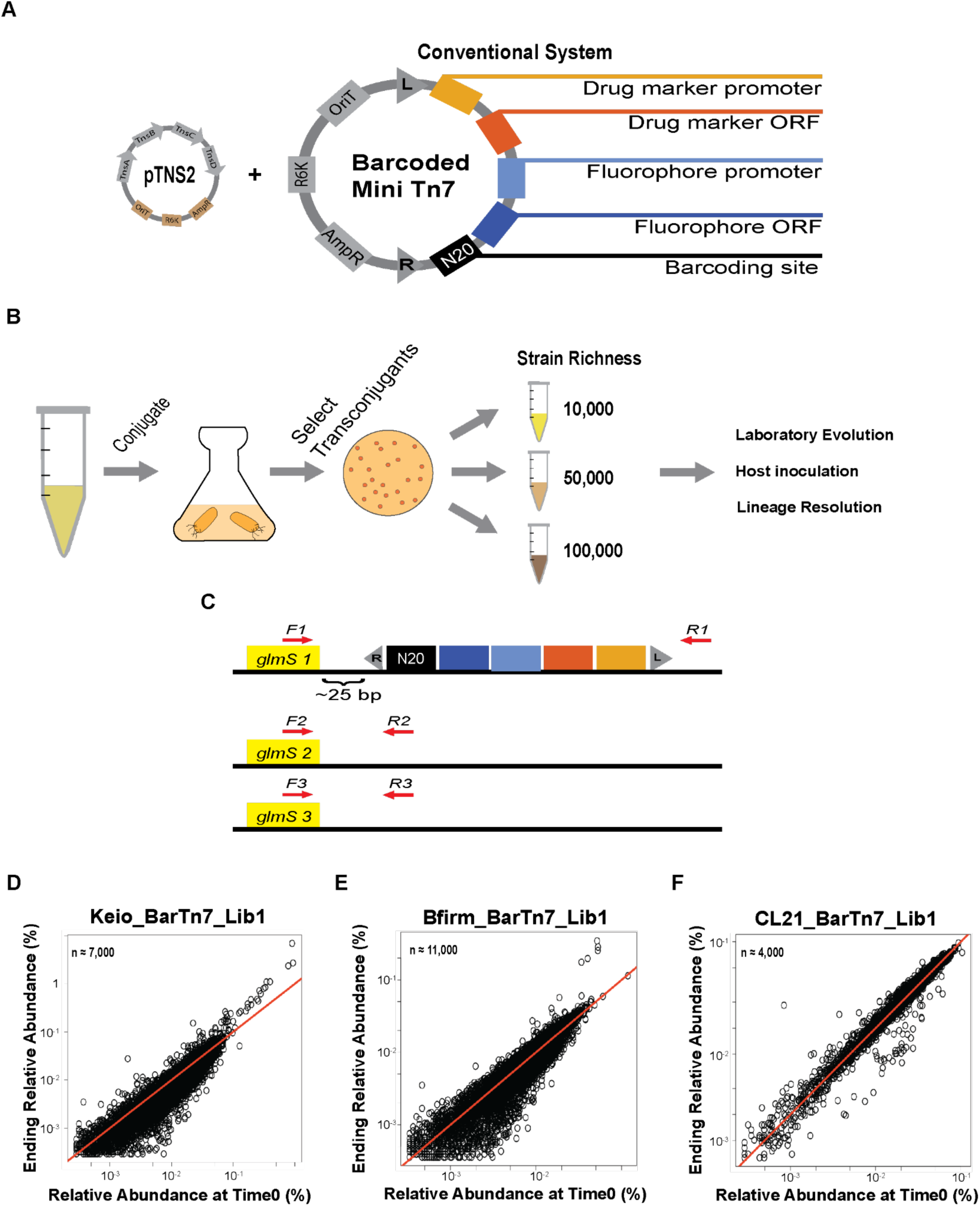
Design and implementation of BarTn7. (**A**) Schematic of the initial BarTn7 vector. Modular genetic cargo are combined onto an R6K-dependent vector. The desired vector is then barcoded and introduced into a desired recipient through tri-parental conjugation including a donor carrying the helper plasmid pTNS2. (**B**) Schematic for barcoding recipient bacteria. In the canonical strategy, two different donor strains are used to deliver the transposase and transposon, carried on two separate plasmids, through conjugation. Transconjugants are pooled to a desired richness appropriate to their experimental use. (**C**) A PCR based strategy for confirming correct localization and orientation of the Tn7 transposon. Species-specific primers are designed to target the junction of the 3’ end of all copies of *glmS* and ∼100 base pairs downstream. For species with multiple homologs of *glmS*, homolog-specific primers are used to confirm Tn7 integrates in monocopy (**D-F**) Results of representative passaging experiments (growth in LB) in BarTn7 populations of three recipient bacteria. In these experiments, we grew each library with serial transfer in LB. The first timepoint (x-axis) represents barcode abundance after an initial outgrowth in 25 mL LB (Time0), while the ending sample (y-axis) is after the third transfer (∼21 population doublings total). The number of detected barcodes in each plot are derived from clustering the starting BarSeq samples using Sheperd (**40**). The red line represents x = y.

We tested at least three single colonies of all 7 recipients to confirm that transposition occurred as expected. In colonies of all the tested recipients, transposition occurred at the expected *attTn7* site 23-25 bp downstream of *glmS*. The species tested ranged from one to three homologs of *glmS.* In species which encode multiple *glmS* homologs (e.g. *P. phytofirmans* PsJN and *P. bryophila* 376MFSha3.1, with 3 and 2 homologs, respectively) Tn7 integrated downstream of a single homolog, confirming previous observations that an initial transposition event attenuates further integration events.

We then generated pooled BarTn7 populations of seven different bacterial species (**Table S1**) of varying richness to use for experimental measurements of population dynamics. For each library, we performed barcode sequencing (BarSeq) to estimate population sizes, ranging from ∼4,000 to ∼280,000 unique strains per library (**Table S1**). In each of two species, *E. coli* BW25113 and *Pseudomonas simiae* WCS417, we generated three populations with log-fold differences in barcode richness (**Table S1**). To test whether individual barcoded strains grow without a biased fitness effect relative to the broader population we passaged each barcoded population individually in LB medium (**Fig. 1D-F, Fig. S1**). While rare strains varied in frequency due to sampling bias during passaging, most strains of all tested species were found at comparable relative abundances between the starting (Time0) and third timepoints, demonstrating that the barcoded lineages were predominantly phenotypically neutral. In populations of the same species the number of rare lineages varying in starting and ending abundances increased in positive correlation with barcode richness due to increasing levels of rare lineages.

### Strain-resolved measurement of switchgrass root colonization

Having confirmed the expected transposition activity of BarTn7 in diverse recipients, we first measured the strain-resolved population dynamics of three root-associated bacterial species during recruitment to the roots of switchgrass plants. We cultivated inoculated switchgrass seedlings with BarTn7 populations of either *Paraburkholderia phytofirmans* PsJN (henceforth “Bfirm”), *Ralstonia* sp. UNC404CL21Col (henceforth “CL21”), or *Pseudomonas simiae* WCS417 (henceforth “WCS417”). For the known plant endophyte *Paraburkholderia phytofirmans* PsJN, we measured colonization of both rhizosphere and root endosphere samples. We also varied the planted medium between high and low phosphate conditions (625 and 30 *µ*M KH_2_PO_4_, respectively) as phosphate deficit is known to increase root biomass, root branching, root hair formation, and root exudate composition in multiple plant species (**15**), all of which could alter the formation of a root microbiome. For Bfirm and CL21, the abundance of each lineage in the rhizosphere generally corresponded with their abundance in the starting community (**Fig. 2A, Fig. S2, Fig. S3**). For WCS417, however, the rhizosphere communities were typically predominated by a subset of the ∼31,000 barcode starting population (**Fig. S2**). Furthermore, for each species, and in most samples, a small number of lineages increased in abundance by 100- to 1,000-fold. Interestingly, some, but not all, of these more successful lineages consistently increased in abundance in multiple replicate rhizosphere samples (**Fig. S3**), potentially indicating pre-existing mutations which increased fitness for rhizosphere colonization. In endosphere samples, consistent with a greater expected bottleneck, the abundance of Bfirm lineages differed dramatically; with most of the initial population lost and the remaining subset representing a greater relative proportion of the resulting community (**Fig. 2B, Fig. S4**). These resulting endosphere communities ranged from 194 to 3,504 barcodes, representing ∼1.8 to ∼31.8% of the starting ∼11,000 barcode library.

**Figure 2.**
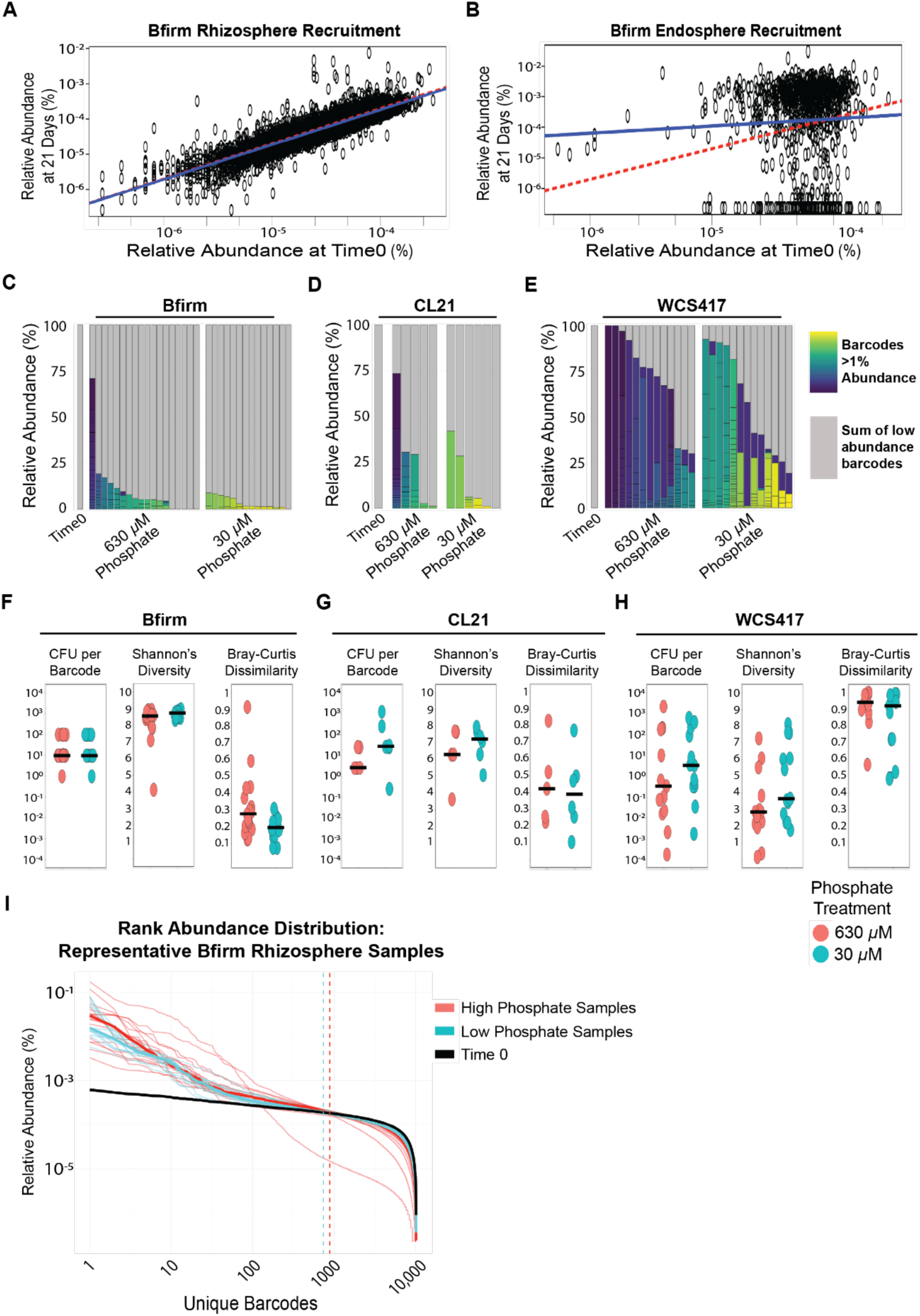
BarTn7 results of switchgrass roots inoculated with barcoded plant-associated bacteria. (**A-B**) Linked rhizosphere and root endosphere samples from the same representative plant inoculated with Bfirm_BarTn7_Lib1 (Bfirm). Solid blue and red dashed lines represent the line of x = y and regressions of barcode relative abundance of the starting population (Time0) versus 21 days post inoculation, respectively. (**C-E**) Proportions of rhizosphere communities derived from barcoded populations of Bfirm_BarTn7_Lib1 (Bfirm), CL21_BarTn7_Lib1 (CL21), and WCS417_BarTn7_Lib2 (WCS417), respectively, present at >1% relative abundance under two different phosphate treatments. Each bar represents a unique rhizosphere sample. Grey portions of each bar represent the sum of the abundances of all barcodes present at < 1% relative abundance and each differently colored portion represents the relative abundance of a unique barcode comprising at least 1% of the community. (**F-H**) Diversity metrics from all rhizosphere samples of Bfirm, CL21, and WCS417 under similar phosphate treatments. Solid lines represent the mean value for each metric. (**I**) Rank abundance distribution of barcodes across the Bfirm BarTn7 population at Time0 versus all rhizosphere samples. Thin lines represent individual samples whereas thicker lines represent the mean across all samples of the same phosphate treatment. Dashed lines represent the point of intersection between experimental samples and Time0. Barcodes are initially ranked along the x axis based on their average abundance across at least four time0 samples.

In the starting populations of each BarTn7 library, no lineages represent more than 1% of the community (**Fig. 2C-E**). In contrast, in the rhizosphere, multiple lineages consistently increased to at least 1% of the community in most experimental samples. These increases differed by species: individual lineages of WCS417 accumulated to greater than 1% of the population in all samples and the proportion of these more abundant barcodes typically represented a greater proportion of the community than for Bfirm or CL21 under both phosphate treatments—in some experiments composing close to 100 percent of the community. Furthermore, the proportion of the community composed by lineages which individually represented more than 1% of the community was consistently higher in high-phosphate samples compared to low-phosphate samples.

To interrogate the cause of single-lineage predominance in the rhizosphere, we tested whether lineages given an early arrival advantage were more competitive than the rest of the population. Briefly, we isolated a uniquely barcoded colony from barcoded CL21 and WCS417 populations and dipped switchgrass roots in saturated cultures of these isolated lineages for 20 minutes prior to transfer to medium inoculated with mixed populations of the corresponding species. For both species, single lineages composed a greater proportion of the rhizosphere communities of plants pre-inoculated with an individual lineage before transfer to the mixed community compared to samples without pre-inoculation (**Fig. S5**). In most cases the isolated lineage represented a significant, if not predominant, proportion of the community. Interestingly, in contrast to plants inoculated with the original mixed inoculation regimen, lineages representing > 1% of the rhizosphere population represented a slightly greater proportion of the population in low-phosphate compared to high-phosphate treated plants, potentially indicating a phosphate-dependent benefit of early arrival to the rhizosphere. In most cases, the preinoculum represented the predominant lineage. Furthermore, for plants inoculated with WCS417, preinoculation was not always sufficient for the pre-inoculum to outperform other members of the mixed community, suggesting additional components required for successful root colonization by an individual lineage of a given species.

BarTn7 additionally enabled the discrimination between two broad drivers of host colonization: colonization events and the proliferation of cells which have already arrived at the root. If the number of barcodes observed represents the minimum number of unique recruitment events, the disparity between unique barcodes and colony forming units (CFU) represents the maximum proportion of the population derived from proliferation of lineages already present in the rhizosphere. For both Bfirm and CL21, the number of CFU recruited per plant was greater than the number of barcodes observed (**Fig. 2F-H**, “CFU per barcode”). However, this ratio changed both by phosphate treatment and species. The ratio of CFU to barcodes was higher in low phosphate samples than high phosphate for both CL21 and WCS417, whereas the ratio was roughly the same for Bfirm. The ratio was also lower for WCS417 under both phosphate treatments than either Bfirm or CL21, likely reflecting either A) the greater starting population richness in the BarTn7 library for this species or B) differences in viability of WCS417 cells when removed from the root and plated on LB agar. Taken together with the number of lineages present at >1% relative abundance in experimental samples, these results highlight potential species- and nutrient-specific root colonization dynamics in WCS417.

To quantify differences in rhizosphere development potentially derived by different means of recruitment, i.e. colonization events versus proliferation, we used conventional ecological measurements to test for differences in barcode evenness and richness in each sample. We first measured Shannon’s diversity, which measures both richness and evenness of taxa observed, in this case of barcoded lineages. In all tested species, high-phosphate samples had lower Shannon’s diversity measurements than their low-phosphate counterparts (**Fig. 2F-H**, “Shannon’s Diversity”), consistent with a smaller number of lineages representing a larger proportion of the community. Consistent with previous measurements, samples inoculated with WCS417 were consistently less diverse than those inoculated with either Bfirm or CL21 despite a larger number of barcodes in the starting population of the former. We then measured Bray-Curtis dissimilarity, which evaluates both richness and abundance of a tested community compared to a separate sample. In this case, experimental samples were compared against the Time0 averages for their respective species, with greater dissimilarity indicating a potential population bottleneck. Consistent with Shannon’s diversity measurements, all species had higher dissimilarity under high-phosphate treatment compared to low-phosphate (**Fig. 2F-H**, “Bray-Curtis Dissimilarity”) and WCS417 samples were more dissimilar from their corresponding Time0 population than either Bfirm or CL21. These results reveal population dynamics common to all three species, e.g. loss of initially rare lineages corresponding to increases in initially abundant counterparts.

Finally, we plotted representative samples against their Time0 averages to draw more specific conclusions from these diversity metrics. In all species, the starting communities are relatively even, with most lineages falling within an order of magnitude of the median (**Fig. 2I, Fig S6**). In contrast, both high- and low-phosphate treated samples revealed subsets of each community either increasing or decreasing in proportion (**Fig. S3, Fig. S6**). The proportion of lineages which changed in abundance relative to their respective Time0 averages was greater for high-phosphate samples compared to those with low-phosphate (**Fig. 2I, Fig. S3, Fig. S6**). Furthermore, and consistent with other metrics, the proportion of the communities composed by these “successful” lineages was also greater under high-phosphate treatment. In all species tested, we observed a trend of initially predominant lineages increasing more in abundance than their rarer counterparts under both phosphate treatments (**Fig. S3, Fig. S6**). Taken together, BarTn7 enabled the measurement of colonization patterns both specific to individual species and dependent on nutrient availability, for example. potential selection for preexisting mutations and competitive exclusion.

### Identification and prediction of adaptive mutations via relative abundance of barcoded lineages

We next tested whether BarTn7 could effectively measure evolutionary dynamics at the strain level. Specifically, we tested whether changes in the relative abundance of barcoded strains could be used to detect the emergence of adaptive mutation(s) in an *in vitro* passaging experiment. We passaged barcoded *E. coli* BW25113 (Keio_BarTn7_Lib4) and *Klebsiella michiganensis* M5a1 (Koxy_BarTn7_Lib5) in liquid LB medium supplemented with increasing concentrations of trimethoprim and measured changes in barcode relative abundance over ∼60 doublings by BarSeq (**Fig. 3A**). After around four passages, we observed unique lineages from each replicate adaptation experiment increase to close to 100% relative abundance (**Fig. 3B, E**). We then plated the final timepoint for four replicate adaptations and screened for colonies matching the barcode of successful lineages by Sanger sequencing.

**Figure 3.**
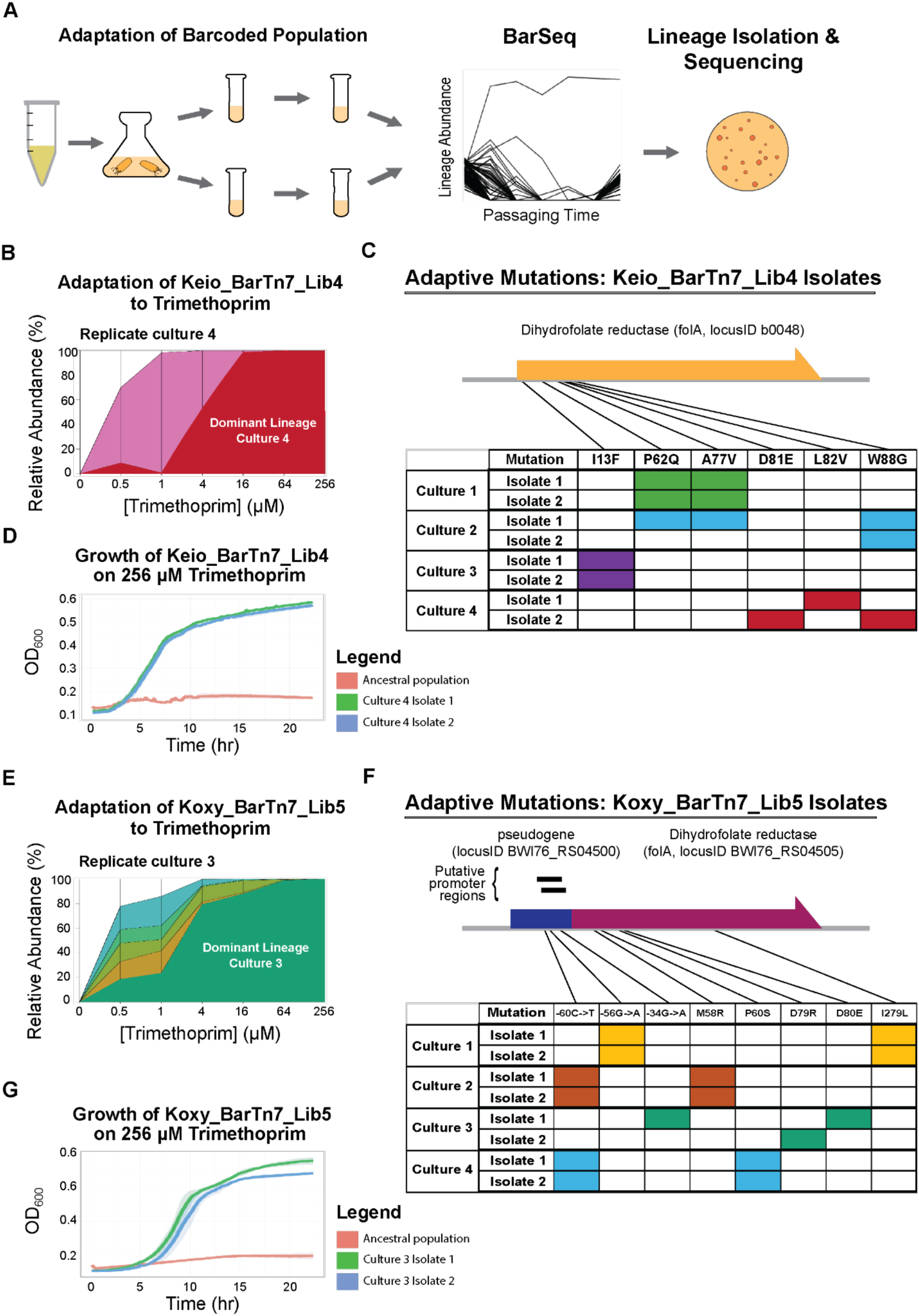
Identification of adaptive mutations during laboratory evolution of barcoded bacterial populations. (**A**) Schematic of experimental design: Keio_BarTn7_Lib4 and Koxy_BarTn7_Lib5 were passaged individually in LB supplemented with increasing concentrations of trimethoprim for ∼60 generations. (**B**) Adaptive trajectories of the most abundant strains at the end of passaging from a representative evolution experiment with Keio_BarTn7_Lib4 and trimethoprim. For the transfer between Time0 and the first trimethoprim-treated sample, the transfer lasted for two nights, shaking at 30°C, whereas all subsequent transfers were overnight at the same temperature. (**C**) Nonsynonymous mutations identified in the *folA* gene of two colonies isolated from the final timepoint of four replicate passages in trimethoprim. Isolates from Culture 4 are the red, dominant lineage in panel **B**. (**D**) Microplate assay comparing the growth of both adapted Keio_BarTn7_Lib4 lineages from Culture 4 and the ancestral population on LB supplemented with 256 µM trimethoprim. (**E**) Adaptive trajectories of the most abundant strains at the end of passaging from a representative set of Koxy_BarTn7_Lib5 samples. For the transfer between Time0 and the first trimethoprim-treated sample, the transfer lasted for two nights with shaking at 30°C, whereas all subsequent transfers were overnight at the same temperature (**F**) Regulatory and nonsynonymous mutations identified in the Koxy *folA* homolog of two replicate colonies isolated from the final timepoint of four replicate passages in trimethoprim. Isolates from Culture 3 are the green, dominant lineage in panel **E**. (**G**) Microplate assay comparing the growth of both adapted Koxy_BarTn7_Lib5 lineages from Culture 3 and the ancestral population on LB supplemented with 256 µM trimethoprim.

After isolating the correct lineages, we first demonstrated with individual growths that these strains were indeed resistant to trimethoprim (**Fig. 3D, G**). Next, we submitted two replicate colonies of four adapted lineages for long-read, whole-genome sequencing to identify causal mutation(s). Using MUMmer4 (**16**), we identified multiple, nonsynonymous single nucleotide polymorphisms in homologs of the *folA* gene for both species (**Fig. 3C, F**), which is known to impact resistance to trimethoprim in both *E. coli* and *Klebsiella* species (**17–18**). Furthermore, at least one such mutation was identified in isolates from separate replicate adaptations of the same library in both species (e.g. Keio_BarTn7_Lib4 *folA* P62Q, A77V, and W88G; or Koxy_BarTn7_Lib5 locusID BWI76_RS04500 −60C->T), emphasizing the impact of these alleles on fitness under treatment with the antibiotic. Interestingly, for Koxy_BarTn7_Lib5, this convergent mutation, as well as two other identified SNPs, occur upstream of *folA* homolog in a predicted pseudogene (locus ID BWI76_RS04500). We therefore reasoned that the observed mutations might be altering the expression of *folA*. In support of this, using Sapphire (**19**) we identified two predicted promoter regions covering all detected mutations.

Lastly, BarTn7 allows us to monitor the population dynamics of low abundance lineages which might otherwise be missed (**Fig. S7**). In all replicate samples strong selective sweeps result in a rapid loss of barcode diversity. Most lineages drop from thousands of reads at early timepoints to near extinction by the middle of the experiment. However, we consistently observe transiently successful lineages competing briefly suggesting partial resistance. Furthermore, especially for Keio_BarTn7_Lib4, many lineages share nearly identical adaptive trajectories early on during passaging, suggesting the simultaneous emergence of resistant strains.

### Comparative sequencing analysis of a multi-species SynCom

We next asked whether BarTn7 could measure ecological dynamics of a population by simultaneously measuring multiple species. We assembled a SynCom from barcoded populations of five species mixed in a 1:1:1:1:1 ratio by OD_600_ (**Fig. 4A**) then outgrew this SynCom in either LB, R2A, or minimal medium supplemented with either 20 mM D-Xylose or D-Glucose as the sole carbon source. We then sequenced the resulting outgrowths via 16S rRNA gene amplification, BarTn7 (BarSeq), and Illumina shotgun whole-genome sequencing (WGS). When comparing the number of reads which aligned to each constituent genome, normalized by genome size, WGS consistently measured each constituent taxon as present in roughly equal relative abundances at the start of the experiment (Time0) (**Fig. 4B**) This trend was consistent across outgrowths in all other media types, with slight discrepancies after growth in LB or RCH2 supplemented with D-xylose (**Fig. 4C-G**). In contrast, even after normalizing by 16S gene copy number, 16S amplicon sequencing differed from WGS by 11.9% on average across all species, samples, and replicates with the exception of one Time0 sample which deviated from the average across all sequencing methods (**Fig. 4G**). Furthermore, the 16S sequences of two SynCom constituents of the genus *Paraburkholderia* were difficult to disambiguate due to their phylogenetic relatedness, a known issue with 16S sequencing. Finally, BarTn7 consistently measured the relative abundance of each species comparably to WGS, differing by an average of 0.9%.

**Figure 4.**
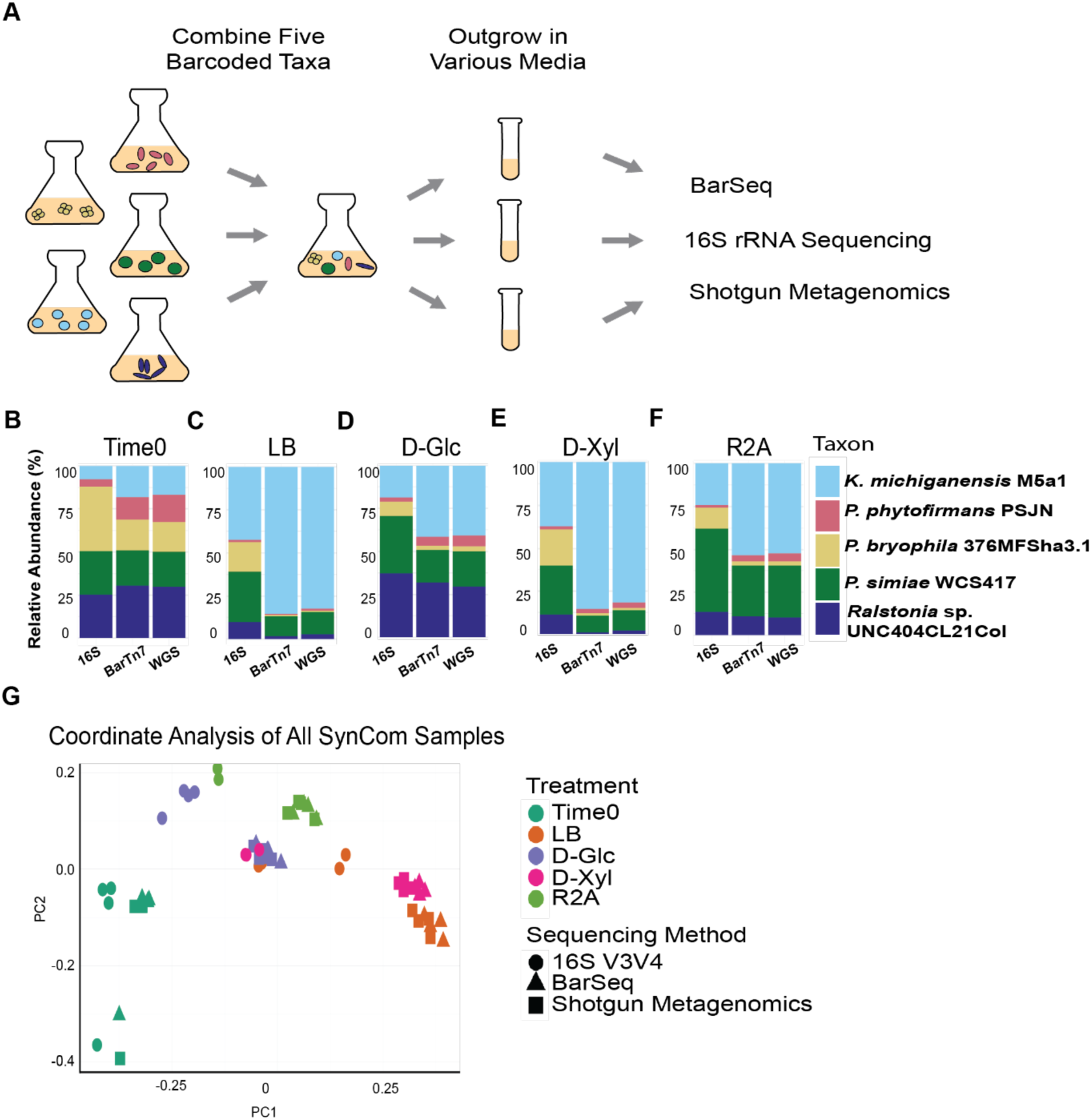
Comparison of a 5-member bacterial synthetic community (SynCom) as measured by three sequencing strategies. (**A**) Experimental schematic: Five barcoded bacterial species were combined at roughly equal OD_600_ into a SynCom, the SynCom was grown in different experimental conditions, and the relative abundance of the community members were measured by BarSeq, 16S rRNA sequencing, and WGS. (**B-F**) Representative plots of the relative abundance of SynCom members as measured by the 3 different methods in the starting population (**B**), and after growth on LB (**C**), RCH2 minimal media supplemented with either D-glucose or D-xylose (**D-E**), or R2A (**F**). (**G**) Principal coordinate analysis of all SynCom samples.

### Optimization of Tn7 delivery vectors for increased efficiency in diverse bacteria

Having demonstrated the utility of the BarTn7 system across multiple experimental designs, we attempted to optimize the system for ease of assembly, efficiency of delivery, and breadth of applicability to various recipient bacteria. First, while deriving and testing various Tn7 vectors, we noticed that some transposon cargo had heterogeneous effects on recipient physiology. Specifically, a barcoded Tn7 vector (pRH54), derived from pRH05 but without the GFP cassette, caused recipient colonies of *E. coli* BW25113 and *K. michiganensis* M5a1 to grow unevenly in size, indicating colony-specific toxicity. We submitted representative colonies from the transconjugation into *E. coli* BW25113 and *K. michiganensis* M5a1 for whole genome sequencing. For *E. coli*, the colonies which grew more quickly had either lost the BarTn7 transposon or never incorporated it to begin with. In contrast, larger *K. michiganensis* M5a1 transconjugants incorporated multiple copies of the Tn7 vector into the *att* site, while the smaller colonies of both contained the Tn7 transposon in the expected location and orientation. As Tn7 is known to cause an increase in recombination at the site of transposition (**20**), we reasoned that if the integration of an engineered Tn7 transposon is indeed toxic, this might select for rare recombination events which alleviate this toxicity. The Tn7 *att* site often sits between two conserved, often essential, operons: the *glmU* operon involved in peptidoglycan synthesis and the *pst* operon involved in phosphate-specific transport. We therefore sought to confirm whether Tn7 integration influenced native gene function by testing four Tn7 delivery vectors (pAD274-pAD277) that contain design elements of previously established Tn7 vectors (see Methods). We conjugated these vectors as before into three recipient bacteria *E. coli* BW25113, *P. simiae* WCS417, and *K. michiganensis* M5a1 to test the impact of varying the sequence sounding the Tn7 cargo on recipient physiology. All tested vectors (pAD274-pAD276) incorporating the Tn7 transposon such that transcription of the cargo was directed towards *glmS*, caused identical, colony-specific growth effects in *E. coli* BW25113 and *K. michiganensis* M5a1. Both of these bacteria have the *pst* operon downstream of *glmS*. In contrast, Tn7 insertions derived from these plasmids showed no toxicity in *P. simiae* WCS417, which contains a nonessential hypothetical gene downstream of *glmS*. Surprisingly, pAD277, which is identical to pAD274 except the reversal of the orientation of the transposon cargo, alleviated this toxicity effect in both *E. coli* BW25113 and *K. michiganensis* M5a1. To eschew any potential negative impact of the Tn7 cargo, we incorporated the transposon cargo of all subsequent plasmids in this reverse orientation.

In most published systems, Tn7 is delivered via a tri-parental conjugation of two plasmids: one encoding the transposon and the other the transposase, into the desired recipient (**12–14**). We reasoned that eliminating the need for the second, “helper” plasmid might increase conjugative efficiency and simplify the generation of BarTn7 libraries in new bacteria. Furthermore, concatenating both elements into a single vector would simplify the optimization of promoters expressing the transposase alongside the transposon cargo via the parts-based strategy of a magic pool system (**9**). We therefore derived two transposon vectors: pRH90 and pRH102. pRH90 is the barcoded version of pAD277, a “two-pot” Tn7 vector containing a kanamycin resistance cassette within the Tn7 transposon, designed to be included in a tri-parental conjugation with a published helper plasmid pTNS2. pRH102 is a single, “one-pot” vector containing the same transposon cargo as pRH90, as well as the Tn7 transposase expressed by the same promoter as in pTNS2. We conjugated these vectors at 1:1:1 or 1:1 ratios of OD_600_, as appropriate, with four different recipient bacteria. With the exception of *P. simiae* WCS417, the “one-pot” delivery system was roughly an order of magnitude more efficient than the two-plasmid delivery method (**Fig. 5A)**. Our results demonstrate that a single BarTn7 plasmid provides an efficient, simplified system for the construction of new libraries in diverse bacteria.

**Figure 5.**
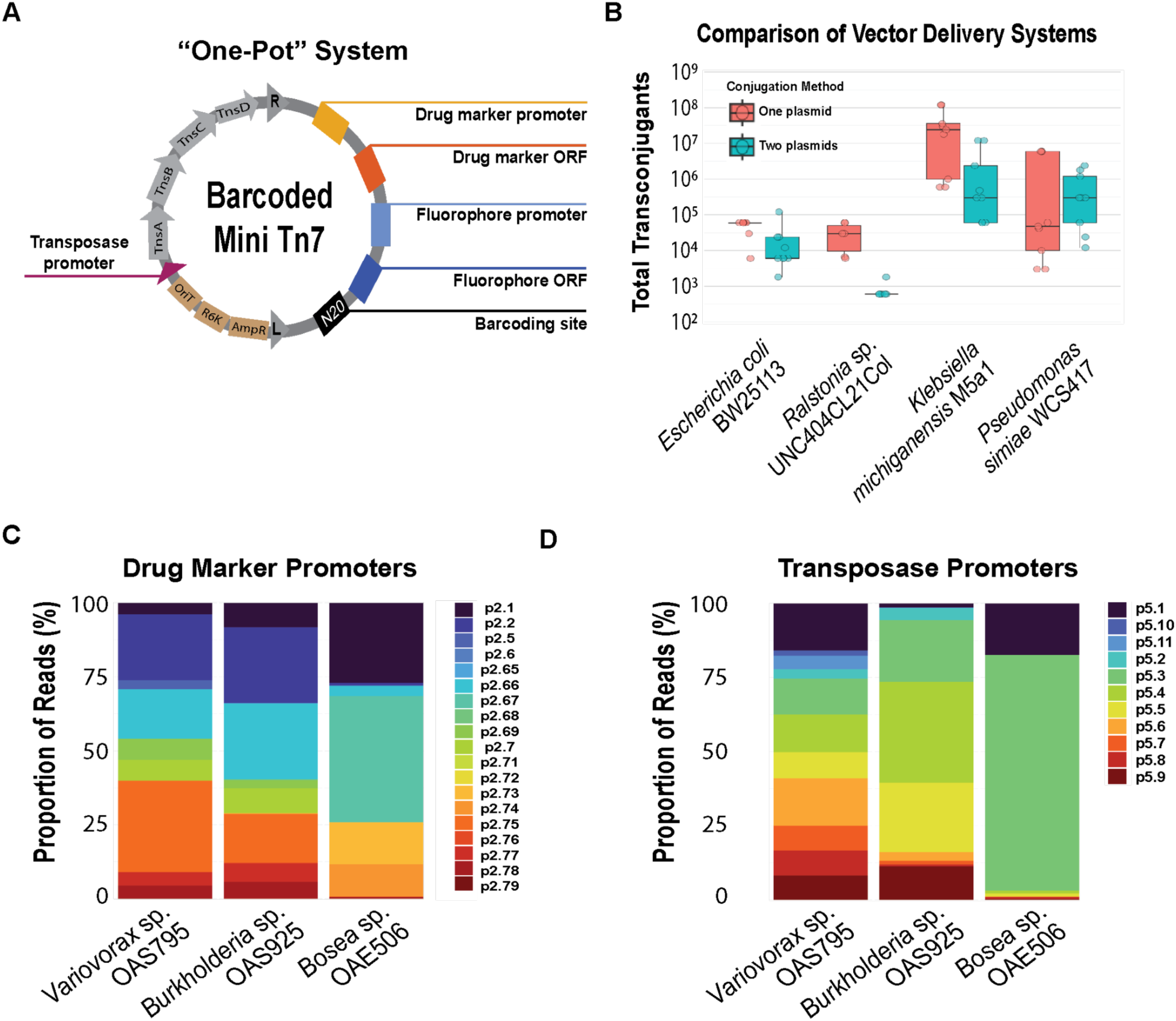
Parts-based optimization of BarTn7 delivery vectors. (**A**) Schematic of the final, single-vector BarTn7 system. Modular genetic cargo, promoters, and *tnsABCD* are combined into a single R6K-dependent vector. The resulting plasmid is then barcoded and introduced into a desired recipient through bi-parental conjugation. (**B**) Comparison of the conjugative efficiency of one (pRH102) and two-pot Tn7 (pRH90) vectors. (**C**) Preference across three species for promoters (part 2 in magic pool design) for kanamycin resistance cassettes. See Table S2 for information about each part. (**D**) Preference across three species for promoters (part 5 in magic pool design) for the Tn7 transposase. See Table S2 for information about each part.

Lastly, we considered the possibility that the application of BarTn7 to new species could be limited by the functionality of the BarTn7 vector parts (promoters, antibiotic resistance genes) in the target bacterium. To streamline this genetic tool development, we implemented the “magic pools” approach for combinatorial vector design and parallel assessment, as described previously for Tn5 and mariner vectors (**9**). For the magic pool designs, we implemented the single-vector system (identical in transposon cargo orientation to pRH102 described in **Figure 5A**). Each BarTn7 magic pool contains 2 different coding sequences for genes conferring resistance to either kanamycin, chloramphenicol, or gentamicin, and a set of promoters for both the transposase and the drug marker in the transposon (**Table S2**). Each vector is linked to its combination of parts by nanopore long read sequencing and a unique barcode in the transposon. Then, by conjugating one of these magic pools into a recipient and performing BarSeq on the pool of resulting transconjugants, we could determine which combination of parts is most efficient for constructing a BarTn7 library

We tested the magic pool conferring kanamycin resistance in three bacterial members of a published SynCom (**21**): *Variovorax* sp. OAS795, *Paraburkholderia sp*. OAS925, and *Bosea sp*. OAE506. For *Variovorax* and *Paraburkholderia*, most of the promoter parts for both the transposase and antibiotic resistance cassette were well-represented (**Fig. 5B-C**). In contrast, the BarTn7 library produced for the *Bosea* recipient was strongly biased towards the *D. vulgaris recA* promoter (p2.67) as the promoter for the drug marker and the *Enterobacter* sp. TBS079 *recA* promoter (p5.4) to express the transposase. Notably, the pTNS2 promoter (p5.11) commonly used for the transposase produced no colonies for either *Paraburkholderia* and *Bosea*, and was poorly represented in the *Variovorax* community. In summary, the resulting magic pools including our changes to transposon orientation and “one-pot” system, serve to optimize the Tn7 system by increasing conjugative efficiency, decreasing potentially toxic effects of transposon integration, and identifying unique promoters and antibiotic resistance cassettes ideal for a variety of recipient bacteria.

## Discussion

This work was designed to increase the resolution of population, evolutionary, and ecological dynamics in microbiome work. We optimized a system (BarTn7) for generating populations of thousands of barcoded lineages, which, apart from the barcode, are isogenic and do not exhibit biased fitness effects relative to one another under standard laboratory growth conditions. These barcoded BarTn7 populations enabled measurements of multiple population dynamics during switchgrass root colonization which would be difficult or impossible to measure with conventional approaches such as 16S rRNA sequencing. Firstly, previous work has demonstrated reduced genetic diversity within the microbiome along a gradient of increasing proximity to the plant (**22–24**) and our experiments confirm this phenomenon at the strain level across the surrounding medium, rhizosphere, and endosphere samples. Secondly, our measurements reveal that these population bottlenecks can be caused by different colonization dynamics. The most relevant of these is the disambiguation of a root-associated population derived from recruitment from the surrounding medium versus the division of cells already in the rhizosphere. Because of the potential for individual lineages being recruited from the surrounding medium multiple times, the disparity between the number of CFU and barcodes observed represents the maximum proportion of the community derived by proliferation, while the number of barcodes represents a minimum number of unique recruitment events. We observed a relatively consistent ratio of CFU to barcodes ranging from 1:1 to 10:1. While there is a potential disparity in cell viability when outgrown in liquid versus on plates (**25**), the fact that this ratio was maintained across all three tested species suggests the ratio observed is accurate. Therefore, we conclude that the proliferation of cells which arrive to the root early on are more impactful to root colonization under our experimental conditions than the continual recruitment of new cells from the medium. Furthermore, this ratio increased under low phosphate conditions for two of three tested bacteria, implying that strain-level root colonization dynamics are dependent on abiotic conditions as is microbiome composition at higher taxonomic levels (**26**).

Moreover, we identified that the importance of early arrival to the root is species-specific. Two species, Bfirm and CL21, colonized the rhizosphere in a relatively neutral fashion: individually-barcoded strains were generally recruited in accordance with their initial predominance in the community. In contrast, WCS417 was observed to colonize the rhizosphere in a biased manner such that individual strains typically came to represent the preponderance of the population. *Pseudomonas* species are known to compete with other bacteria and fungi in the rhizosphere (**27–28**). Indeed, mutualistic species of *Pseudomonas* have been shown to inhibit root colonization by closely-related *Pseudomonas* pathogens (**29**) leading us to question the possibility of strain-level competitive interactions during the development of a root-associated WCS417 population. Our pre-inoculation experiments show that both WCS417 and CL21 strains are likely to outcompete their surrounding counterparts when given an early advantage during colonization. Importantly, some of the predominance of these single lineages can be attributed to a greater starting load. However, the pre-inoculum is unlikely to be sufficient to cause the predominance of one strain in a population of hundreds of thousands of cells. Furthermore, the pre-inoculum did not always predominate, suggesting other population dynamics at play which contribute to colonization. These experiments therefore show potential competitive exclusion, a mechanism conferring historical contingency (**30**) which, to our knowledge, has not been observed at such a high taxonomic resolution.

BarTn7 also enables measurements of ecological dynamics in bacterial communities. We show that measurement of DNA barcodes as a proxy for strain abundance in a SynCom is more accurate than 16S sequencing. This is likely due to several factors: 16S sequencing is known to be impacted by primer bias and the 16S rRNA gene can be encoded in multiple copies, the number of which varies by species, while Tn7 is chromosomally integrated in monocopy. We also show that BarTn7 is almost identically accurate to shotgun metagenomics, the gold standard for interpreting community composition. Furthermore, because BarTn7 requires a targeted BarSeq PCR amplicon and only 50 base-pair reads, it is less expensive than both 16S and whole genome sequencing.

Finally, BarTn7 enables and hastens the measurement of evolutionary dynamics in bacterial populations. Adaptive mutations are easily identifiable by tracking barcode frequencies. Unlike metagenomic sequencing, the knowledge of a specific barcode associated with greater fitness relative to the ancestral population enables the linkage of a potential mutant allele to a specific strain. Indeed, we unintentionally identified several lineages which appear to have preexisting mutations which confer increased fitness in our root inoculation experiments. Thus, evolutionary experiments can be performed without laborious screening of colonies for phenotypes and mutant alleles of interest, and at a fraction of the cost.

Others have used various forms of DNA barcoding, both with Tn7 and other genetic systems, for tracking closely related lineages (**31–34**). In contrast to existing methods, our magic pool system allows a variety of genetic parts to be rapidly screened to identify an ideal combination for a given recipient. Indeed, many of the promoters we used in our magic pool were more efficient in expressing the Tn7 transposase than that from the published pTNS2 helper plasmid. These new parts combinations enabled transposition of Tn7 into recipient strains which were previously recalcitrant. Next, combining the Tn7 transposon and transposase into a single plasmid increased conjugative efficiency in three out of four tested recipient species. Finally, during the development of our vectors we identified previously undescribed, recipient-specific toxicity of certain Tn7 transposon vector designs. Alterations in the orientation of the transposon cargo serve to alleviate this toxicity and maintain the fitness neutrality of barcoded recipient strains. In combination, BarTn7 enables the rapid implementation of inexpensive, high-resolution lineage tracking in diverse and new strains of bacteria.

## Materials & Methods

### Bacterial strains and cultivation

Bacterial strains (**Table S3**) were maintained in glycerol stocks stored at −80°C. Media used were lysogeny broth (LB; 10 g/L tryptone, 5 g/L yeast extract, and 5 g/L NaCL), R2A (0.5 g/L casein hydrolysate, 0.5 g/L D-glucose, 0.5 g/L soluble starch, 0.5 g/L yeast extract, 0.3 g/L dipotassium phosphate, 0.3 g/L sodium pyruvate, 0.25 g/L casein peptone, 0.25 g/L meat peptone, 0.024 g/L magnesium sulfate), or RCH2 (100 mLs 10X salts, 10 mLs Wolfe’s Vitamins, 10 mLs Wolfe’s Minerals, 50 mLs 0.6M PIPES buffer, 820 mLs water). Liquid precultures were inoculated with single colonies or 1 mL aliquots of barcoded libraries in 5 mL and 50 mL LB plus antibiotic and/or diaminopimelic acid (DAP), respectively, and grown at 30°C shaking at 180 RPM. Optical density was measured at 600 nm.

### Construction of BarTn7plasmids

For a full list of plasmids used in this study, along with details about their construction, see **Table S4**. Primers and gBlocks used for construction are listed in **Table S5** and **Table S6**. To construct a mini-Tn7 vector backbone, two DNA fragments were synthesized as gBlocks from IDT: gRH1, containing an ampicillin resistance cassette and the Tn7 right arm and gRH2, containing the Tn7 left arm and a barcoding region containing an internal pair each of BbsI and BsmBI cut sites. A DNA fragment containing an R6K conditional origin of replication and an origin of transfer was amplified from the pTNS2 “helper” plasmid (Addgene #64968) (**35**) with oRH1 and oRH2. This PCR product, gRH1, and gRH2 were ligated by Gibson assembly, yielding pRH01. An antibiotic resistance cassette and fluorophore were added as transposon cargo by Golden Gate assembly with pRH01, pHLL216 (Kan promoter), pHLL238 (KanR gene), pTKO5 (GFP promoter), and pTKO16 (GFP gene) using BbsI to yield pRH02. Random, 20 nucleotide DNA barcodes were generated and incorporated by Golden Gate assembly with BsmBI into pRH02 to yield pRH05, as previously described (**9**). All restriction enzymes were purchased from Thermo Fisher Scientific. After each Golden Gate assembly, the resulting vector was purified using a Zymo DNA Clean and Concentrator Kit (Cat. #D4004), re-digested with the appropriate restriction enzyme for 16 h to remove residual background vector and purified again with the Zymo kit. During the course of the project, we found via whole plasmid sequencing that our pRH05 construct contained two copies of both the kanamycin resistance gene and GFP, thus libraries made with this barcoded Tn7 library contain a larger cargo region within the Tn7 arms. All other barcoded plasmid libraries we made in this study were confirmed with whole plasmid sequencing to only have a single copy of the cargo genes. We constructed a subsequent barcoded Tn7 vector (pRH54) where we substituted the GFP promoter/gene with a transcriptional terminator part (pAD213), but found Tn7 insertions derived from this construct to be toxic to some recipient bacteria.

To explore different BarTn7 vector designs, particularly with respect to the sequences between the Tn7 arms and antibiotic resistance cassette, along with the orientation of transcription of the antibiotic resistance gene, we constructed a set of 4 non-barcoded Tn7 vectors; pAD274 contains sequences including FRT sites derived from pUC18T-mini-Tn7T_Gm (**35**), pAD275 contains sequence features from pTn7-SCOUT12.G (**36**), pAD276 has sequences from Tn7 integration vector (**12**), and pAD277 is identical to pAD274 except the orientation of the antibiotic resistance cassette is flipped. We found that Tn7 insertions derived from pAD277 were not toxic in any recipient bacteria. Subsequently, we incorporated the DNA barcodes into pAD277 as described (**9**) to generate pRH90. We constructed pRH102, a one-pot BarTn7 vector that combines elements of pRH90 and the pTNS helper plasmid, via Golden Gate assembly of plasmids pAD278, pHLL216, pHLL238, pAD281, pAD213, and pAD279; followed by barcode incorporation with an additional round of Golden Gate assembly.

### Preparation of barcoded bacterial communities

Barcoded plasmid libraries were electroporated into a DAP auxotrophic *E. coli* WM3064 conjugative donor strain and grown overnight at 37°C in 50 mL LB with carbenicillin and DAP. The resulting donor library was split into 1 mL aliquots in 25% glycerol and stored at −80°C. For each conjugation, an entire aliquot of the transposon donor library, the wild-type recipient, and, when necessary for two-vector systems, a second WM3064 donor carrying either pTNS2 or pTNS3, were outgrown overnight. Each strain was washed 3X in LB without antibiotics, mixed 1:1:1 by OD_600_, plated on recipient medium plus DAP, and incubated at 30°C. The next day, the conjugation mix was scraped and re-plated on recipient medium plus antibiotics, without DAP. The conjugation media and selection media was LB for all libraries except for *Paraburkholderia* sp. OAS925, *Variovorax* sp. OAS795, and *Bosea* sp. OAE506. For these 3 bacteria, we used R2A.

Integration of the Tn7 transposon was confirmed by PCR with primers annealing to the 3’ end of the recipient-specific *glmS* gene and roughly 100 bp downstream of *glmS*. For recipient strains with multiple *glmS* homologs, each homolog was tested with unique primer pairs. After validating Tn7 insertion and localization, barcoded communities were pooled by plating the conjugation mixture as above and scraping the desired number of colonies into 50 mL recipient medium plus antibiotic and the number of uniquely barcoded strains was confirmed by BarSeq. We made multiple glycerol stock aliquots for each library for future experiments.

### *In vitro* growth assays of barcoded communities

Aliquots of barcoded libraries were outgrown overnight in 50 mL rich medium plus 100 µg/mL kanamycin. For passaging assays to demonstrate the libraries were predominantly phenotypically neutral, libraries were initially cultured in 50 mL LB plus kanamycin until saturation, then diluted back into fresh 5 mL LB cultures plus kanamycin to repeat the process, for a total of 3 passages. Relative abundances of barcoded lineages were compared between the initial outgrowth (Time0) and the third passage. For *in vitro* adaptation experiments, barcoded communities were first outgrown in 50 mL LB supplemented with 100 µL/mL kanamycin. The following day, these cultures were diluted to a starting OD_600_ of 0.01 in quadruplicate, 5 mL cultures in fresh LB with 0.5 µM trimethoprim. After 1-2 days of growth (2 days for first passage in the minimal trimethoprim concentration and 1 day for all subsequent concentrations), 1 mL samples were collected for BarSeq to represent one passage, and each culture was diluted 1:1000 in fresh LB medium with a slightly higher trimethoprim concentration. For co-inoculation of multiple bacterial species, barcoded pools of each species were grown, individually and overnight, in 50 mL LB plus 100 µg/mL kanamycin, washed to remove antibiotics, and combined at equal ODs immediately prior to use. Quadruplicate Time0 samples were taken from cultures of both the individual species and the SynCom. The SynCom was then inoculated to a starting OD_600_ of 0.01 in quadruplicate, 5 mL volumes of LB, R2A, or RCH2 with 20 mM D-glucose or D-xylose as a sole carbon source and grown overnight before sampling.

### Plant growth conditions and root inoculation

Seeds of *Panicum virgatum* v. Blackwell (switchgrass) were a generous gift from Dr. Jenny Mortimer. Seeds were surface sterilized by a 1-minute wash in 70% (v/v) ethanol, a 15-minute wash in 5% (v/v) sodium hypochlorite, and three rinses with sterile water. Seeds were stratified for four days in the dark at 4°C on ½ Murashige and Skoog agar (Cassion labs MSP01, 1 g/L MES hydrate, 0.9% w/v BactoAgar) before germination under a 12:12 day light:dark cycle at 24°C. At the emergence of the first set of true leaves (4-5 d) individual seedlings were transferred to 80 mL ½ MS with 0.3% BactoAgar in magenta boxes (Fisher Scientific). High or low phosphate treatments (625 and 30 µM, respectively) were performed by preparing soft agar with appropriate amounts of MS basal salts with or without phosphate (Cassion labs MSP11). Bacteria were grown in LB overnight with 100 µg/mL kanamycin and washed without antibiotic prior to inoculation at a concentration of 10^5^ CFU per mL of soft agar.

After three weeks of incubation, rhizosphere fractions were collected by removing the roots and vortexing for 30 s in a 5 mL final volume of LB with antibiotic as appropriate. The washed roots were removed and weighed. To collect endosphere fractions, roots were surface-sterilized by vortexing 30 s in 70% ethanol, 30 s in 5% sodium hypochlorite plus 0.01% (v/v) Triton-X 100, then washed three times in sterile water. One mL of the final wash solution was plated on LB to assess surface sterility. Surface-sterilized roots were homogenized in 1 mL LB plus antibiotic in a Qiagen TissueLyser II at 30 Hz for fifteen seconds. All fractions were outgrown in LB plus kanamycin overnight prior to gDNA extraction.

### DNA sequencing, barcode quality filtration, and statistical analysis

Shotgun whole-genome metagenomics of the BarTn7 SynCom was performed by Novogene on an Illumina NovaSeq instrument. V3V4 amplicon sequencing of the 16SrRNA gene was also performed by Novogene. Reads were first trimmed for quality control using Trimmomatic (**37**) then aligned to either the reference genomes or 16S sequences of the SynCom constituents. Species relative abundance was assessed by comparing the proportion of reads which aligned to reference sequences associated with each of the five species, normalized by number of 16S homologs or genome size for 16S or whole genome sequencing, respectively. BarSeq was performed as described in Wetmore, *et al* (**38**), except we used dual-indexed primers to identify instances of index hopping. For the adaptive evolution experiments on trimethoprim, we performed Nanopore long-read whole genome sequencing (Plasmidsaurus) and mutations were identified by comparing the *folA* sequences from the ancestral and adapted strains (NCBI Taxon IDs 511145 and 290337) using Geneious Prime (**39**).

Quadruplicate Time0 samples of barcoded communities were collected prior to all experiments. DNA from frozen pellets was isolated using the Qiagen DNeasy Blood and Tissue kit according to the manufacturer’s instructions. DNA barcodes were filtered by those with Phred quality scores >30 for each base. For single timepoint experiments, barcodes were clustered using Shepherd (**40**), then filtered by removing barcodes with fewer than three reads in experimental samples. For timecourse experiments, barcodes were clustered using Shepherd, then analyzed using the union of all barcodes across corresponding Time0 samples. This method allowed for the possibility of low-abundance barcodes which increase in proportion over the course of the experiment. For SynCom experiments, sets of barcodes were determined for each species from their individual Time0 samples, then used to determine the species of lineages observed in SynCom samples. Strains which could not be unambiguously assigned to a single species were never more than 5% of these samples. In instances wherein low numbers of barcodes came to predominate the community, the error rate during clustering was loosened to account for a greater frequency of sequencing errors relative to the predominant barcode sequence.

### BarTn7 magic pool construction

The BarTn7 magic pool design and cloning approach were largely modeled from those used in the original random barcode transposon site sequencing (RB_TnSeq) *mariner* and Tn5 vectors (**9**). The BarTn7 design is a modular 6 or 7 part Golden Gate assembly: Part_1 contains Tn7 *tns* enzymes and the right Tn7 arm and is invariable, Part_2 contains the promoter for the antibiotic resistance marker and varies in the magic pool, Part_3 is the antibiotic resistance gene and is also a variable part in the pool (both Part_2 and Part_3 are identical to the original parts designed for Tn5 and *mariner*, and some of these part vectors are used in this study), Part_4 contains the DNA barcodes, the Tn7 left arm, oriT, the ampicillin resistance gene, and the R6K origin of replication, Part_5 is the promoter for the Tn7 enzymes and is a variable part, Part_6 is a promoter for the fluorescence gene, while Part_7 is the fluorescence gene coding sequence. Part_6 and Part_7 are also within the transposon and can be subbed out for any cargo of interest. In the magic pools designed and used in this study, we used a short combined Part_6-Part_7 sequence, and this part was invariable. All parts were cloned into a universal holding vector (pJW52) as previously described (**9**). Detailed information about all plasmid constructions can be found in **Table S4**. We constructed 3 magic pools in this study: pAD286 contains 19 Part_2 promoter variants, 2 Part_3 variants encoding proteins that confer resistance to kanamycin, and 11 variants of the Part_5 promoter. Promoter parts were derived from several “housekeeping” genes of various bacteria. pAD287 and pAD288 are identical to pAD286 except we substituted the Part_3 variants with those that conferred resistance to gentamicin and chloramphenicol (see **Table S2** for full designs). After Golden Gate assembly with BbsI to generate the magic pool, we transformed into EC100D pir+ cells, and selected for colonies on LB plates with carbenicillin. We pooled together ∼10,000 colonies per magic pool and extracted plasmid DNA from the library. To link each DNA barcode to its variable parts (Part_2, Part_3, and Part_5) in the plasmid, we used long-read nanopore sequencing. For pAD286, we could confidently link 6,004 unique DNA barcodes to at least one variable part. We could do the same for 9,435 unique barcodes in pAD287 and 6,183 unique barcodes in pAD288. We then transformed the magic pool plasmids into the WM3064 conjugation strain for delivery into recipient bacteria.

## Acknowledgements

The authors thank Tiffany Lowe-Power, Johan Leveau, Eduardo Ruiz, and members of the Deutschbauer lab for invaluable discussions and input. This material by m-CAFEs Microbial Community Analysis & Functional Evaluation in Soils, a Science Focus Area led by Lawrence Berkeley National Laboratory, is based upon work supported by the US Department of Energy, Office of Science, Office of Biological & Environmental Research under contract no.: DE-AC02-05CH11231.

---

**Table S1:** BarTn7 Libraries.

| Strain ID | Parental strain | BarTn7 library | Estimated library size | Construction note |
| --- | --- | --- | --- | --- |
| AMD1351 | *Escherichia coli* BW25113 | Keio BarTn7 Lib1 | 7,000 | made via conjugation with AMD1386 (*E. coli* WM3064 carrying pRH05 barcoded transposon) |
| AMD1352 | *Escherichia coli* BW25113 | Keio BarTn7 Lib2 | 90,000 | made via conjugation with AMD1386 (*E. coli* WM3064 carrying pRH05 barcoded transposon) |
| AMD1353 | *Escherichia coli* BW25113 | Keio BarTn7 Lib3 | 280,000 | made via conjugation with AMD1386 (*E. coli* WM3064 carrying pRH05 barcoded transposon) |
| AMD3525 | *Escherichia coli* BW25113 | Keio BarTn7 Lib4 | 385,000 | made via conjugation with AMD3509 (*E. coli* WM3064 carrying pRH90 barcoded transposon) |
| AMD1346 | *Ralstonia sp.* UNC404CL21Col | CL21 BarTn7 Lib1 | 4,000 | made via conjugation with AMD1386 (*E. coli* WM3064 carrying pRH05 barcoded transposon) |
| AMD1348 | *Pseudomonas simiae* WCS417 | WCS417 BarTn7 Lib1 | 5,000 | made via conjugation with AMD1386 (*E. coli* WM3064 carrying pRH05 barcoded transposon) |
| AMD1349 | *Pseudomonas simiae* WCS417 | WCS417 BarTn7 Lib2 | 31,000 | made via conjugation with AMD1386 (*E. coli* WM3064 carrying pRH05 barcoded transposon) |
| AMD1350 | *Pseudomonas simiae* WCS417 | WCS417 BarTn7 Lib3 | 110,000 | made via conjugation with AMD1386 (*E. coli* WM3064 carrying pRH05 barcoded transposon) |
| AMD1843 | *Klebsiella michiganensis* M5a1 | Koxy BarTn7 Lib1 | 72,000 | made via conjugation with AMD1386 (*E. coli* WM3064 carrying pRH05 barcoded transposon) |
| AMD3526 | *Klebsiella michiganensis* M5a1 | Koxy BarTn7 Lib5 | 347,000 | made via conjugation with AMD3509 (*E. coli* WM3064 carrying pRH90 barcoded transposon) |
| AMD1844 | *Paraburkholderia bryophila* 376MFSHa3.1 | Burk376 BarTn7 Lib1 | 45,000 | made via conjugation with AMD1386 (*E. coli* WM3064 carrying pRH05 barcoded transposon) |
| AMD1846 | *Paraburkholderia phytofirmans* PsJN | BFirm BarTn7 Lib1 | 11,000 | made via conjugation with AMD1386 (*E. coli* WM3064 carrying pRH05 barcoded transposon) |
| AMD1847 | *Pseudomonas fluorescens* SBW25 | SBW25 BarTn7 Lib1 | 62,000 | made via conjugation with AMD1386 (*E. coli* WM3064 carrying pRH05 barcoded transposon) |

**Table S2:**
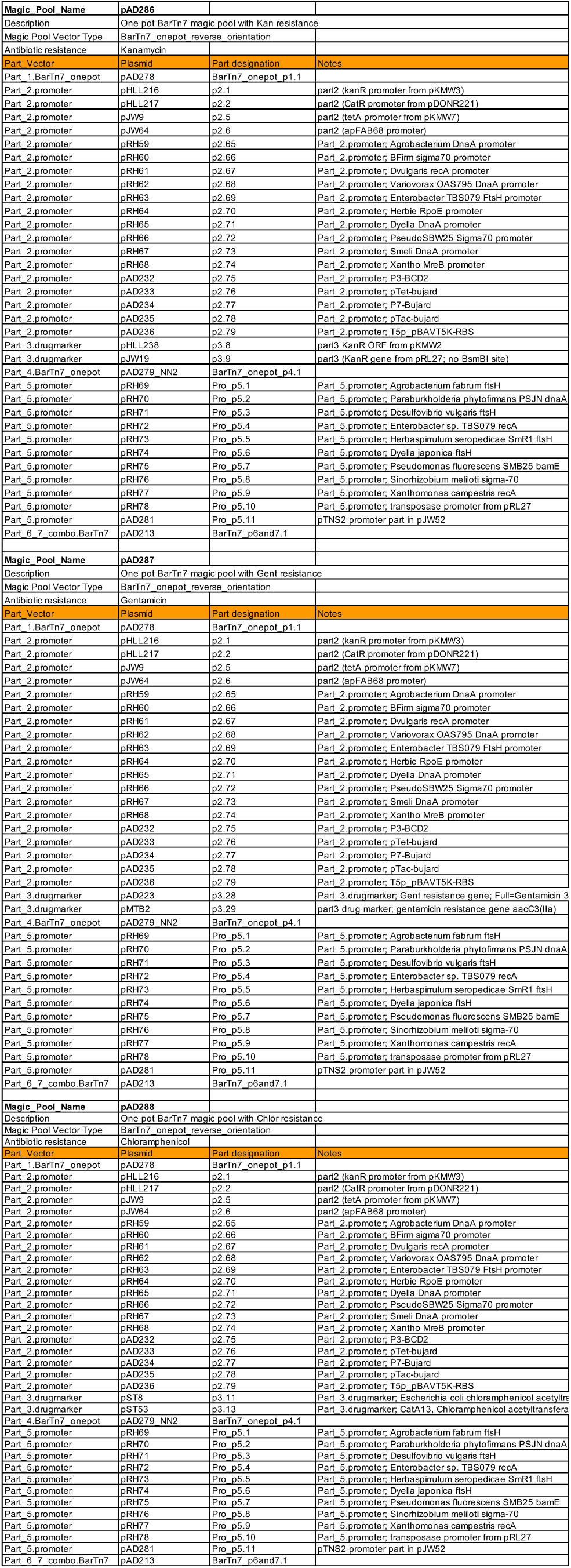
Magic Pool Parts.

**Table S3:**
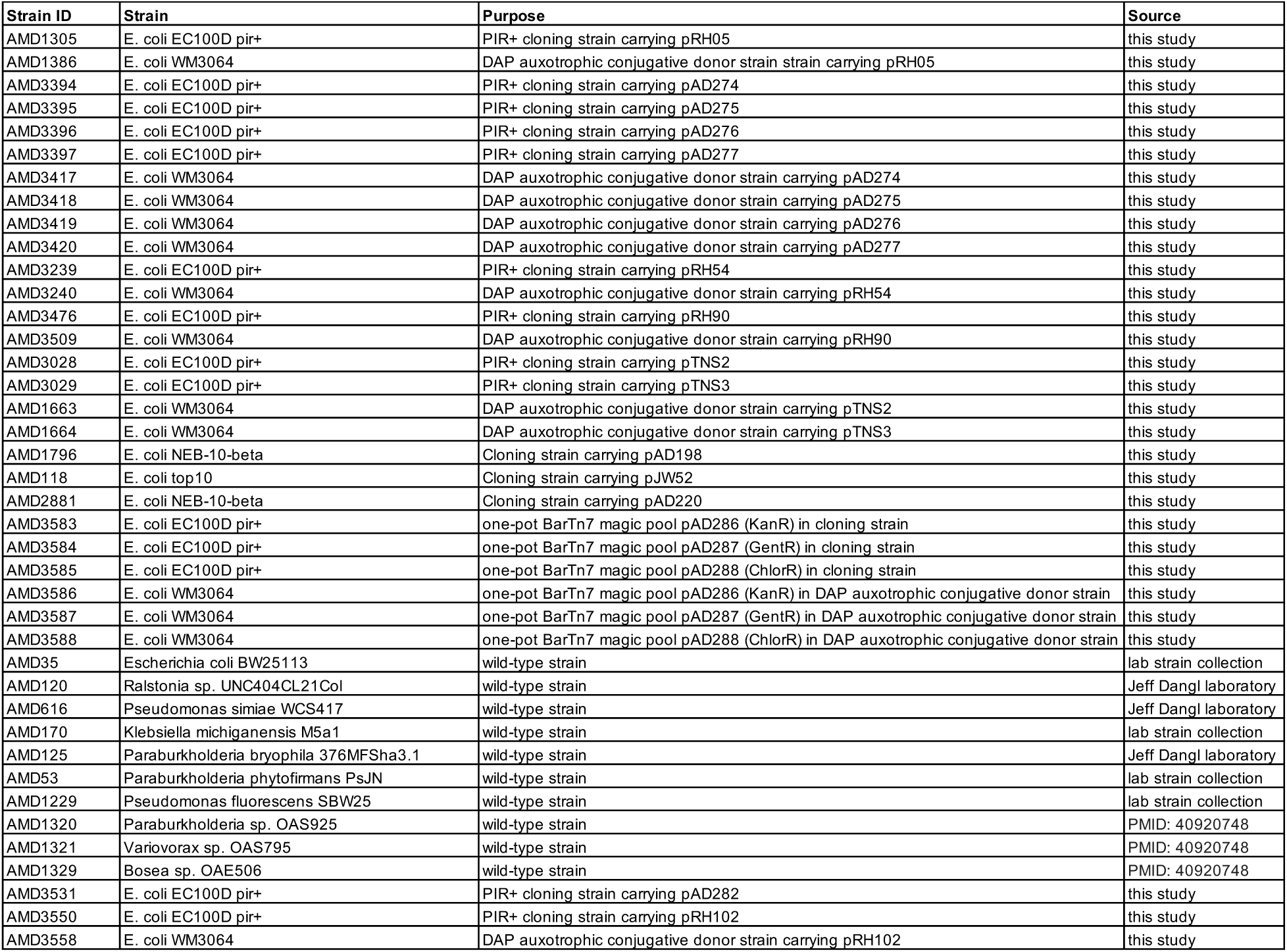
Strains Used.

**Table S4:**
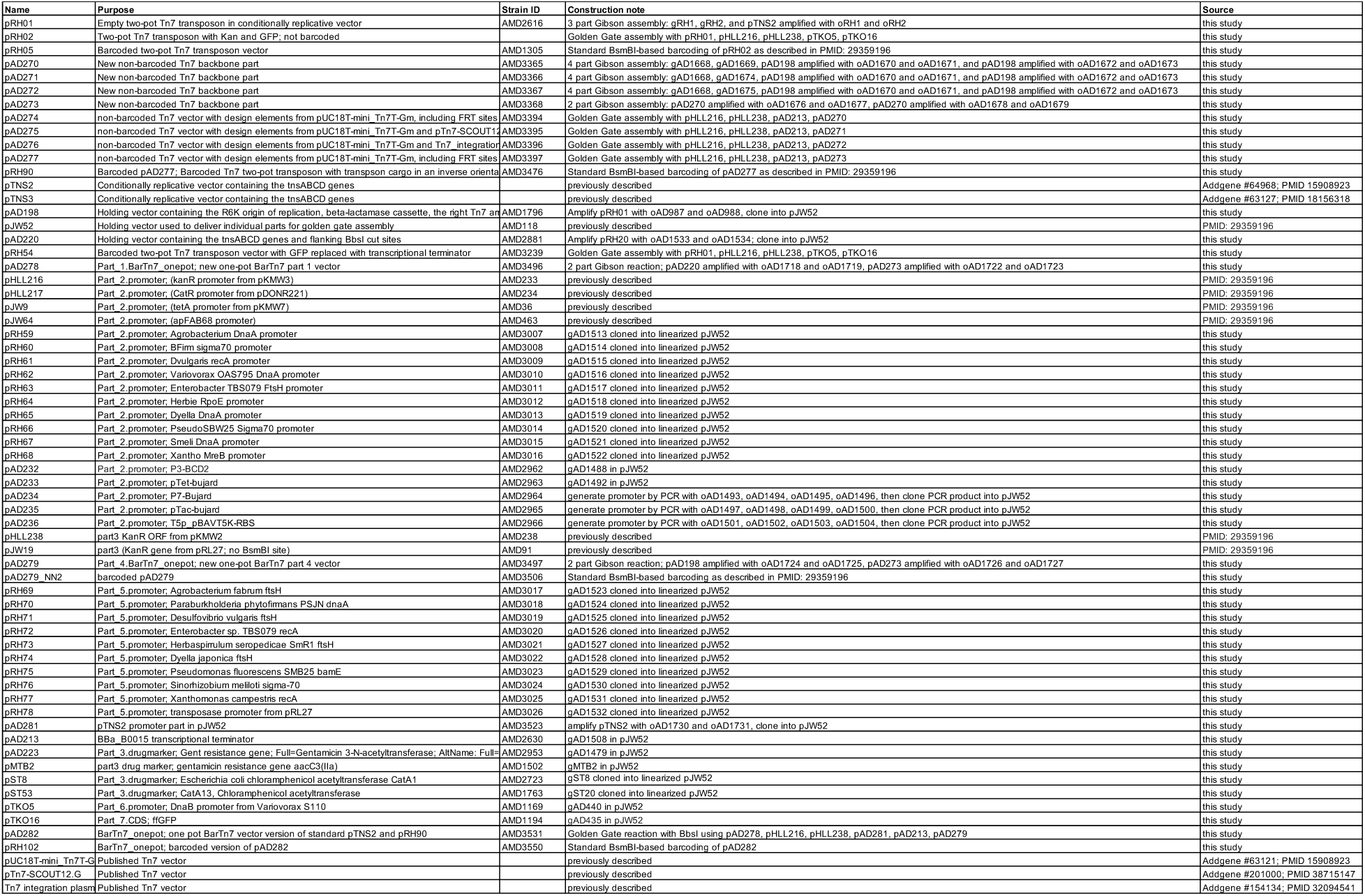
Plasmids Used.

**Table S5:**
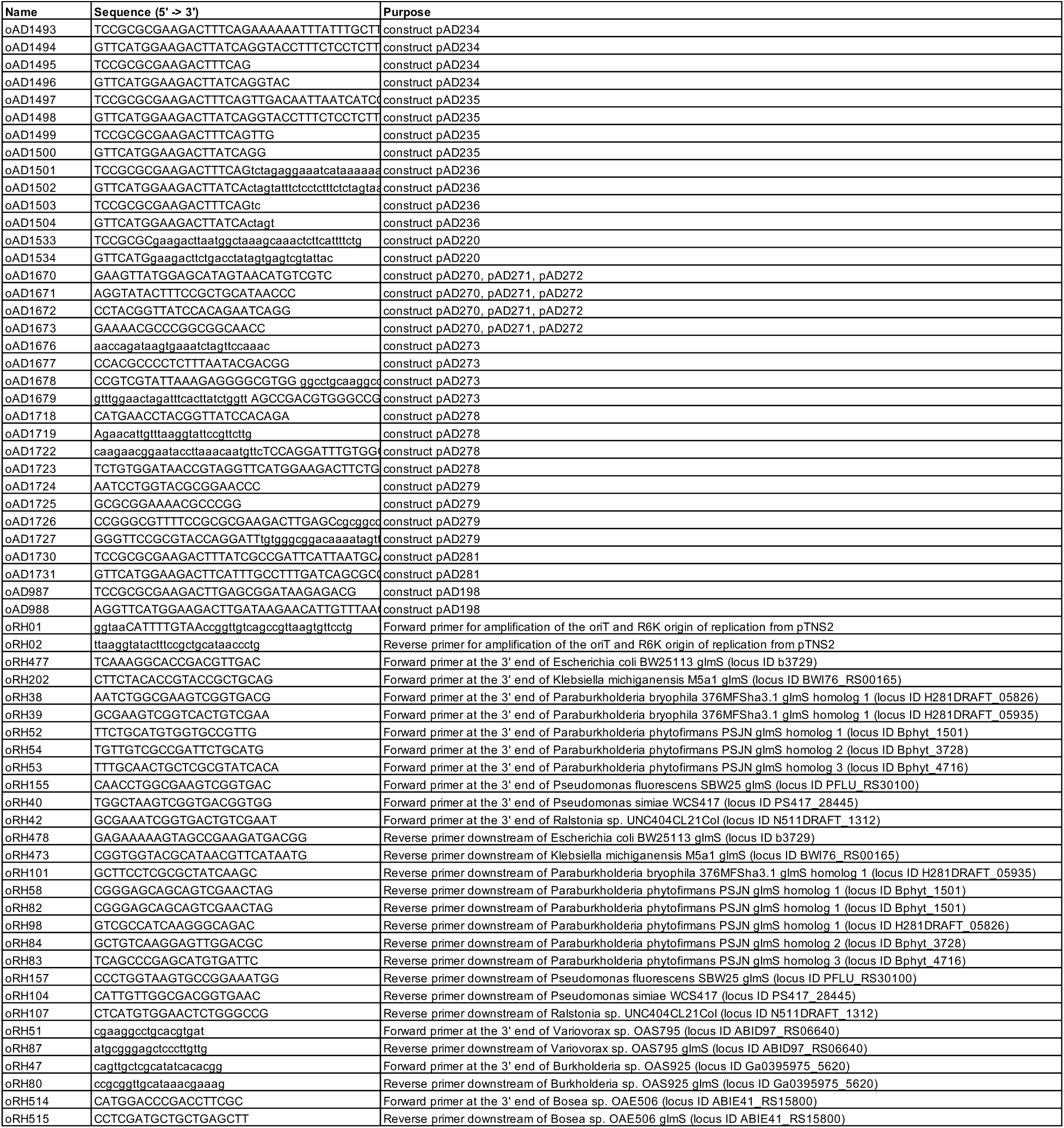
Primers Used.

**Table S6:**
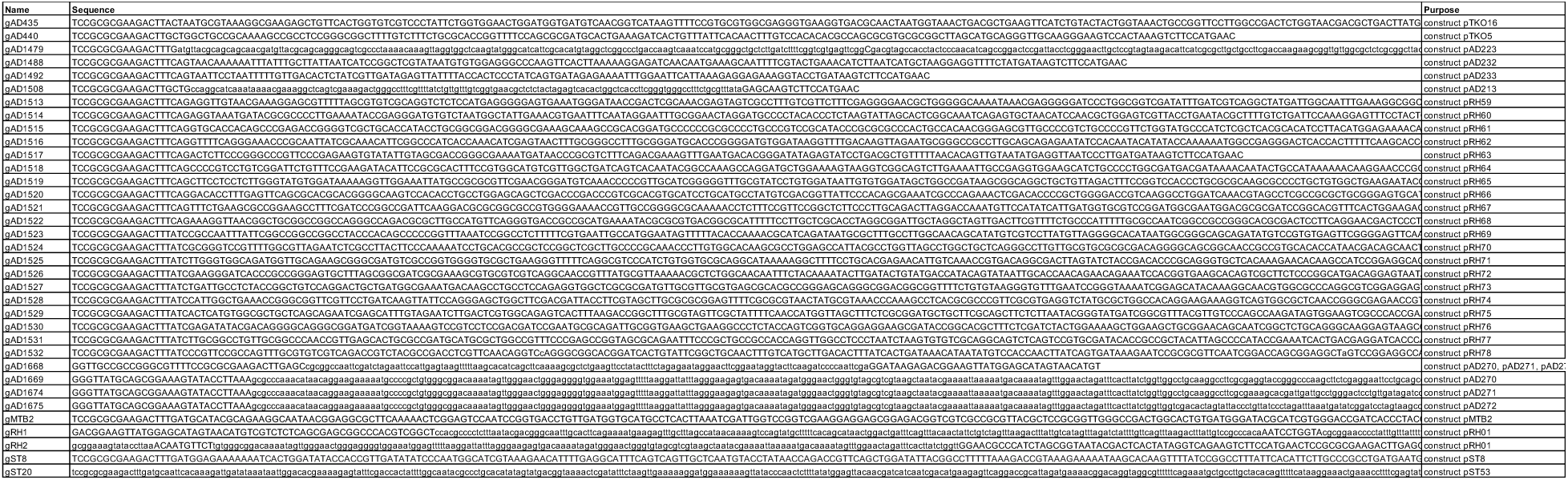
gBlocks Used.

