## Supplemental Figures for "BarTn7: Optimizing Bacterial Lineage Tracking at Sub-Species Resolution for Population Dynamics in Ecological and Evolutionary Studies"

#### Figure S1

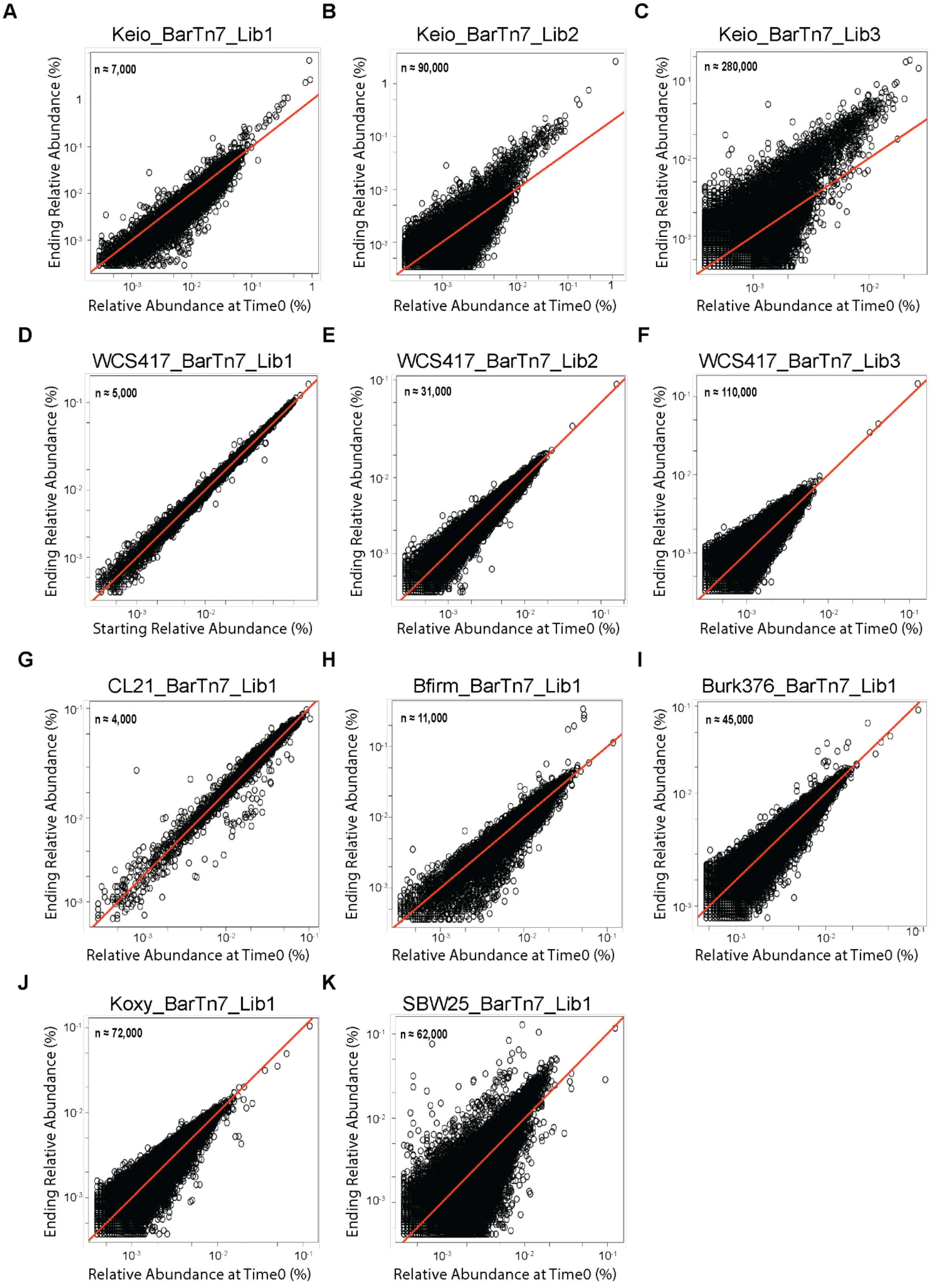

A
Keio_BarTn7_Lib1
n ≈ 7,000
Ending Relative Abundance (%)
Relative Abundance at Time0 (%)
B
Keio_BarTn7_Lib2
n ≈ 90,000
Ending Relative Abundance (%)
Relative Abundance at Time0 (%)
C
Keio_BarTn7_Lib3
n ≈ 280,000
Ending Relative Abundance (%)
Relative Abundance at Time0 (%)
D
WCS417_BarTn7_Lib1
n ≈ 5,000
Ending Relative Abundance (%)
Starting Relative Abundance (%)
E
WCS417_BarTn7_Lib2
n ≈ 31,000
Ending Relative Abundance (%)
Relative Abundance at Time0 (%)
F
WCS417_BarTn7_Lib3
n ≈ 110,000
Ending Relative Abundance (%)
Relative Abundance at Time0 (%)
G
CL21_BarTn7_Lib1
n ≈ 4,000
Ending Relative Abundance (%)
Relative Abundance at Time0 (%)
H
Bfirm_BarTn7_Lib1
n ≈ 11,000
Ending Relative Abundance (%)
Relative Abundance at Time0 (%)
I
Burk376_BarTn7_Lib1
n ≈ 45,000
Ending Relative Abundance (%)
Relative Abundance at Time0 (%)
J
Koxy_BarTn7_Lib1
n ≈ 72,000
Ending Relative Abundance (%)
Relative Abundance at Time0 (%)
K
SBW25_BarTn7_Lib1
n ≈ 62,000
Ending Relative Abundance (%)
Relative Abundance at Time0 (%)

**Figure S1**. Results of passaging experiment in barcoded populations of all seven tested recipient bacteria. In these experiments, we grew each library with serial transfer in LB. The ending sample (y-axis) is after the third transfer (~14 population doublings total). The number of detected barcodes in each plot are derived from clustering the starting BarSeq samples using Sheperd (**40**). (**A-C**) Passaging experiments for three different barcoded populations of *E. coli* BW25113 varying in barcode richness (Keio). (**D-F**) Passaging experiments for three different barcoded populations of *P. simiae* WCS417 (WCS417) varying in barcode richness. (**G-K**) Passaging experiments in the remaining tested species: *Ralstonia* sp. UNC404CL21Col (CL21), *P. phytofirmans* PSJN (Bfirm), *Paraburkholderia bryophila* 376MFSha3.1 (Burk376), *Klebsiella michiganensis* M5al (Koxy), and *Pseudomonas fluorescens* SBW25 (SBW25), respectively. Plots for Keio_BarTn7_Lib1, CL21_BarTn7_Lib1, and Bfirm_BarTn7_Lib1 are repeated from Figure 1 for comparison.

#### Figure S2

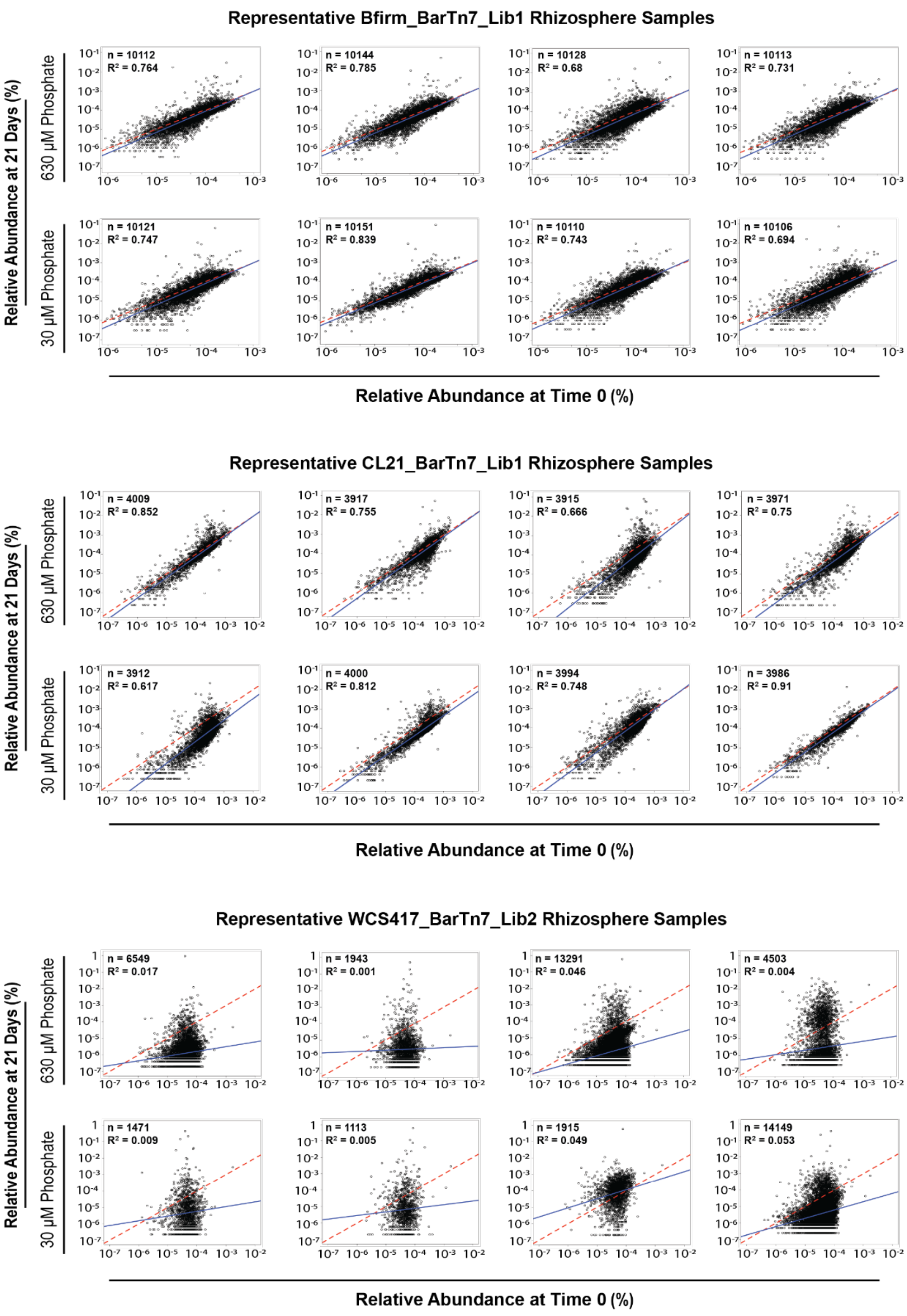

Representative Bfirm_BarTn7_Lib1 Rhizosphere Samples
Relative Abundance at 21 Days (%)
630 µM Phosphate
30 µM Phosphate
n = 10112
R² = 0.764
n = 10144
R² = 0.785
n = 10128
R² = 0.68
n = 10113
R² = 0.731
n = 10121
R² = 0.747
n = 10151
R² = 0.839
n = 10110
R² = 0.743
n = 10106
R² = 0.694
Relative Abundance at Time 0 (%)
Representative CL21_BarTn7_Lib1 Rhizosphere Samples
Relative Abundance at 21 Days (%)
630 µM Phosphate
30 µM Phosphate
n = 4009
R² = 0.852
n = 3917
R² = 0.755
n = 3915
R² = 0.666
n = 3971
R² = 0.75
n = 3912
R² = 0.617
n = 4000
R² = 0.812
n = 3994
R² = 0.748
n = 3986
R² = 0.91
Relative Abundance at Time 0 (%)
Representative WCS417_BarTn7_Lib2 Rhizosphere Samples
Relative Abundance at 21 Days (%)
630 µM Phosphate
30 µM Phosphate
n = 6549
R² = 0.017
n = 1943
R² = 0.001
n = 13291
R² = 0.046
n = 4503
R² = 0.004
n = 1471
R² = 0.009
n = 1113
R² = 0.005
n = 1915
R² = 0.049
n = 14149
R² = 0.053
Relative Abundance at Time 0 (%)

**Figure S2**. Barcode starting and ending abundances in representative rhizosphere samples. All plots represent data derived from independent plants. Samples are separated by high- and low-phosphate treatment and by inoculation with barcoded populations of Bfirm, CL21, or WCS417. Solid blue and red dashed lines represent the line of x = y and regressions of barcode relative abundance at Time0 versus 21 days post inoculation, respectively. The number of barcodes observed at sampling (n) and $R^2$ values for each linear regression are listed in the top left of each plot.

**Figure S3**

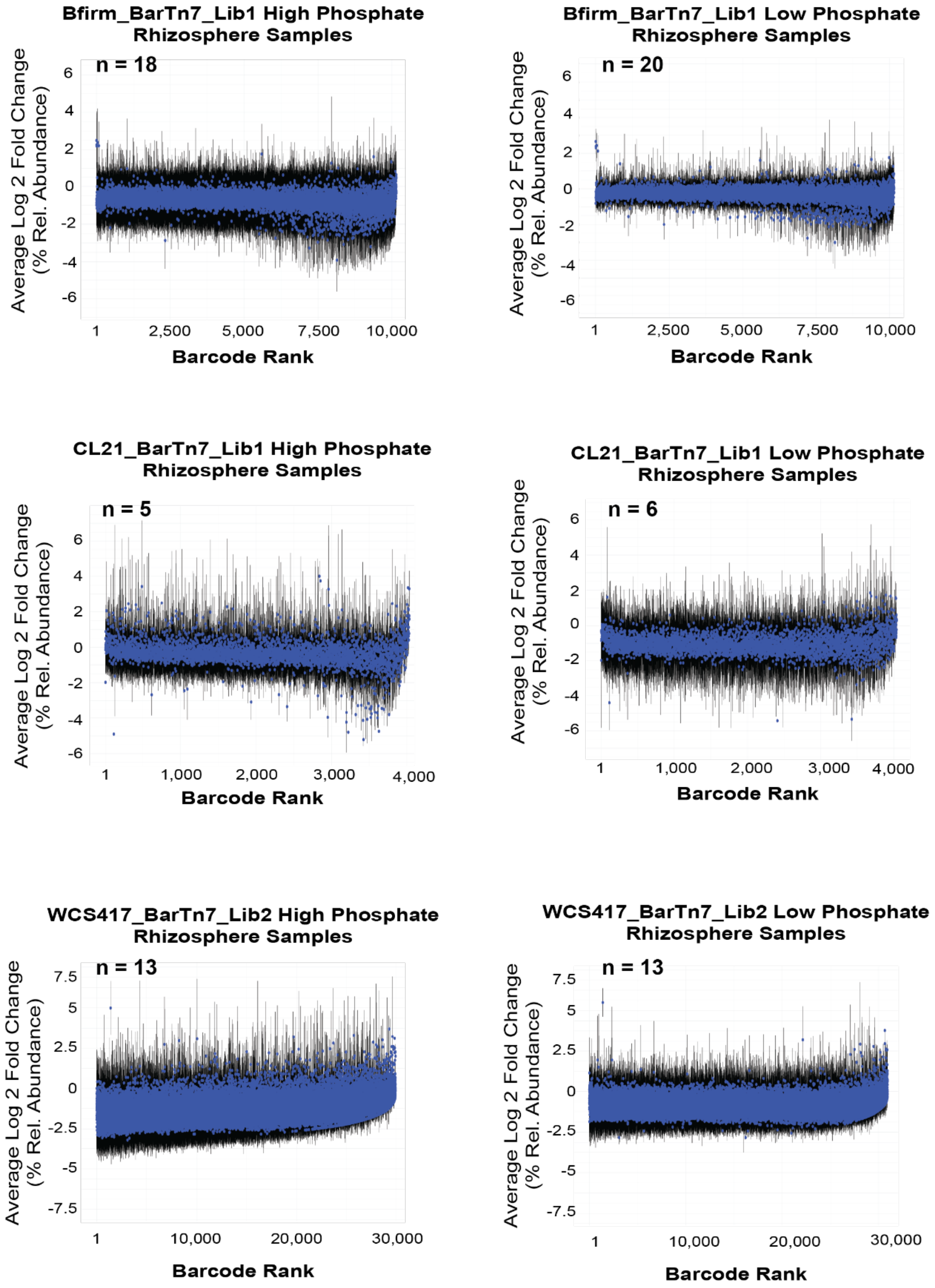

Bfirm_BarTn7_Lib1 High Phosphate Rhizosphere Samples
n = 18
Average Log 2 Fold Change (% Rel. Abundance)
6
4
2
0
-2
-4
-6
1
2,500
5,000
7,500
10,000
Barcode Rank
Bfirm_BarTn7_Lib1 Low Phosphate Rhizosphere Samples
n = 20
Average Log 2 Fold Change (% Rel. Abundance)
6
4
2
0
-2
-4
-6
1
2,500
5,000
7,500
10,000
Barcode Rank
CL21_BarTn7_Lib1 High Phosphate Rhizosphere Samples
n = 5
Average Log 2 Fold Change (% Rel. Abundance)
6
4
2
0
-2
-4
-6
1
1,000
2,000
3,000
4,000
Barcode Rank
CL21_BarTn7_Lib1 Low Phosphate Rhizosphere Samples
n = 6
Average Log 2 Fold Change (% Rel. Abundance)
6
4
2
0
-2
-4
-6
1
1,000
2,000
3,000
4,000
Barcode Rank
WCS417_BarTn7_Lib2 High Phosphate Rhizosphere Samples
n = 13
Average Log 2 Fold Change (% Rel. Abundance)
7.5
5
2.5
0
-2.5
-5
-7.5
1
10,000
20,000
30,000
Barcode Rank
WCS417_BarTn7_Lib2 Low Phosphate Rhizosphere Samples
n = 13
Average Log 2 Fold Change (% Rel. Abundance)
7.5
5
2.5
0
-2.5
-5
-7.5
1
10,000
20,000
30,000
Barcode Rank

**Figure S3**. Average log2 fold change in barcode relative abundance. Fold changes in rhizosphere abundance relative to the starting population (Time0), separated by inocula BarTn7 library and phosphate treatment. Blue dots represent the mean log2 fold change in abundance for each barcode across all experimental samples. Barcodes are ranked from left to right based on their average relative abundance at Time0 in ascending order. Black vertical lines represent the standard deviation of the mean for each barcode.

**Figure S4**

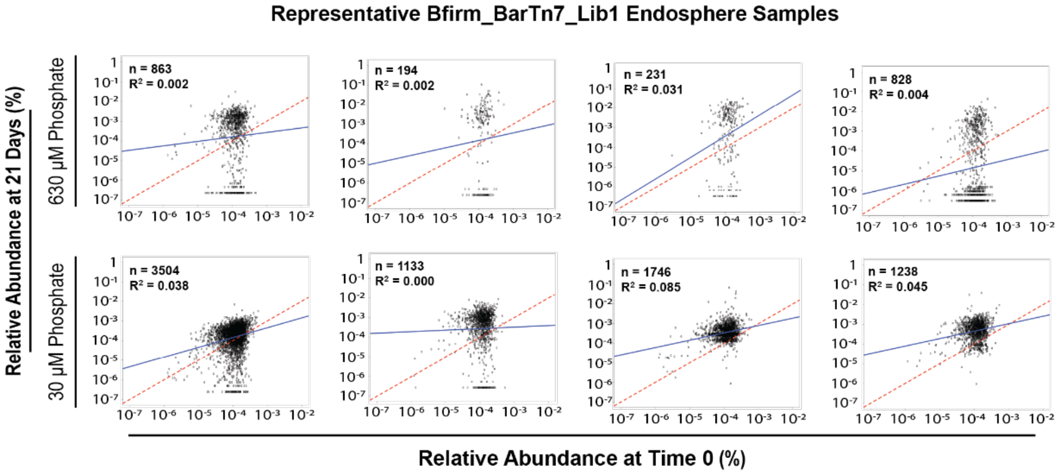


**Figure S4**. Barcode starting and ending abundances in representative root endosphere samples from plants inoculated with Bfirm_BarTn7_Lib1. Each plot represents an independent plant experiment. Samples are separated by high- and low-phosphate treatment. Solid blue and red dashed lines represent the line of x = y and regressions of barcode relative abundance at Time0 versus 21 days post inoculation, respectively. Number of barcodes observed at sampling (n) and $R^2$ values for each linear regression are listed in the top left of each plot.

**Figure S5**

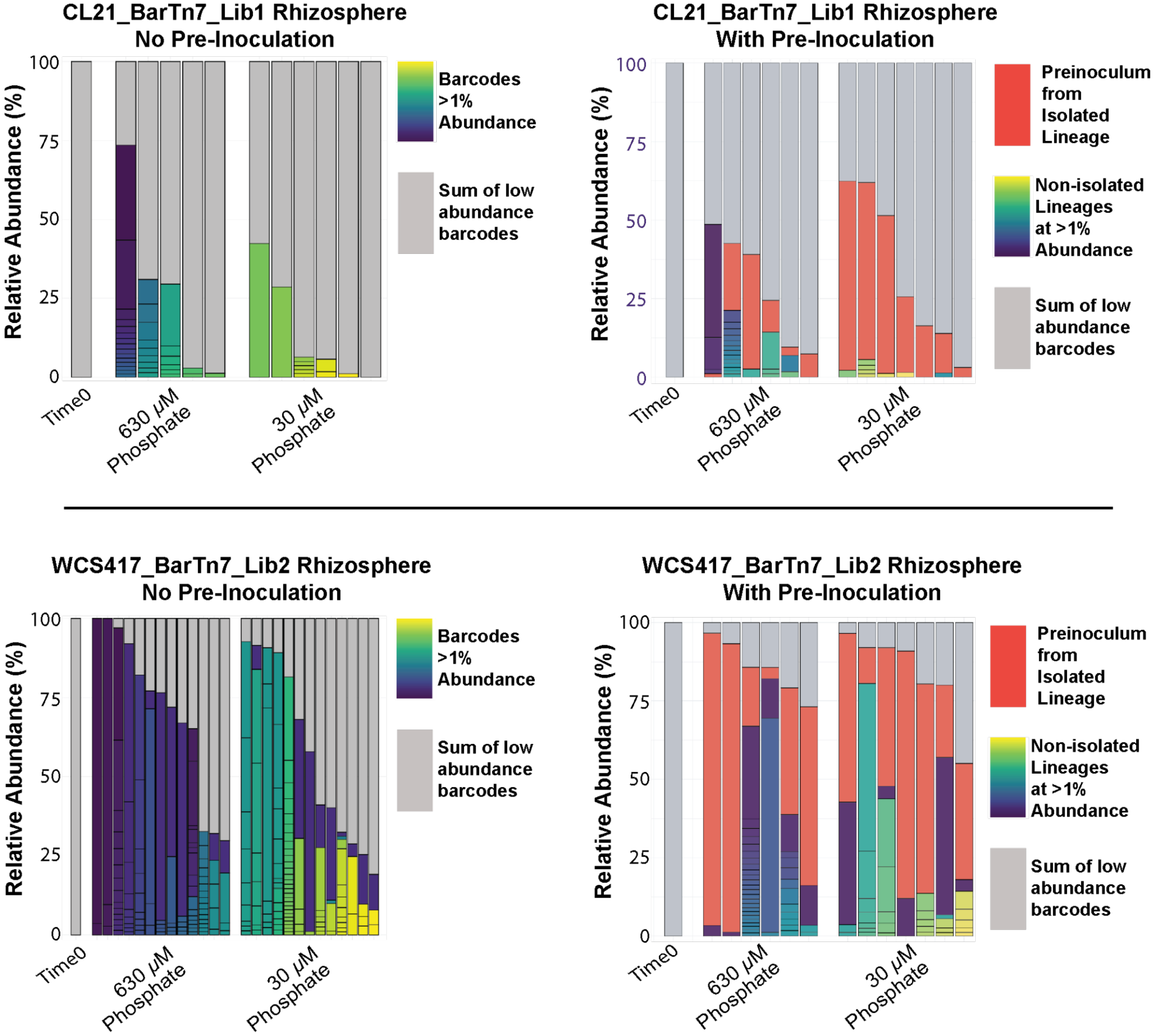


**Figure S5**. Proportions of rhizosphere communities present at >1% relative abundance in mixed-inoculation and pre-inoculation experiments. Grey bars represent the sum of the abundances of all barcodes present at <1%. Lineages representing ≥1% of the community, but not used as pre-inoculum for pre-inoculation experiments, are depicted by colored bars in viridis. The isolated lineage used for pre-inoculum in each library is colored consistently in red across experiments. The no pre-inoculation data from **Figure 2D-E** are shown for comparison.

### Figure S6

A

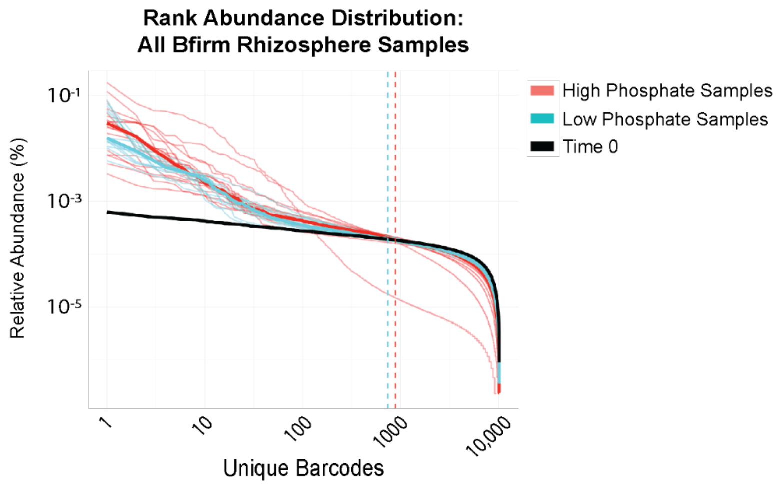


B

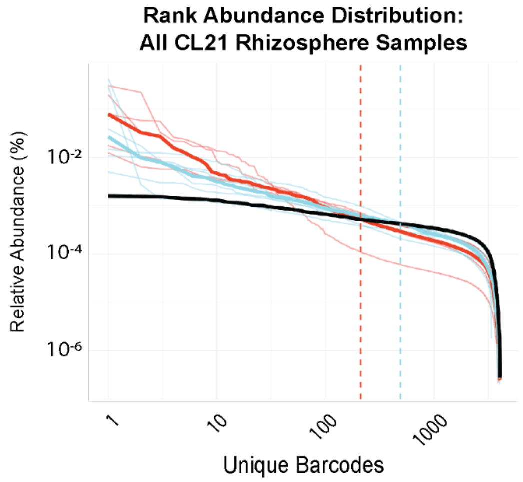


C

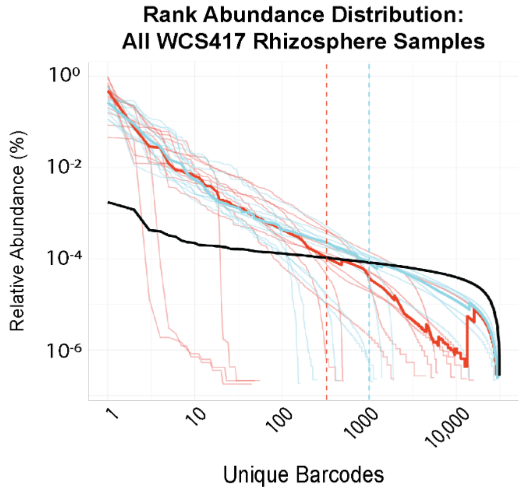

**Figure S6**. Rank abundance distribution of barcodes across all tested root inoculation samples. Ranked abundance of barcodes from three tested libraries, Bfirm_BarTn7_Lib1 (Bfirm), CL21_BarTn7_Lib1 (CL21), and WCS417_BarTn7_Lib2 (WCS417), are plotted at Time0 versus all rhizosphere samples per library. Thin lines represent individual samples whereas thicker lines represent the mean across all samples of the same phosphate treatment. Dashed lines represent the point of intersection between experimental samples and Time0. Barcodes are initially ranked along the x-axis based on their average abundance across at least four Time0 samples. Data from Bfirm rhizosphere samples are copied from Figure 2I for comparison.

#### Figure S7

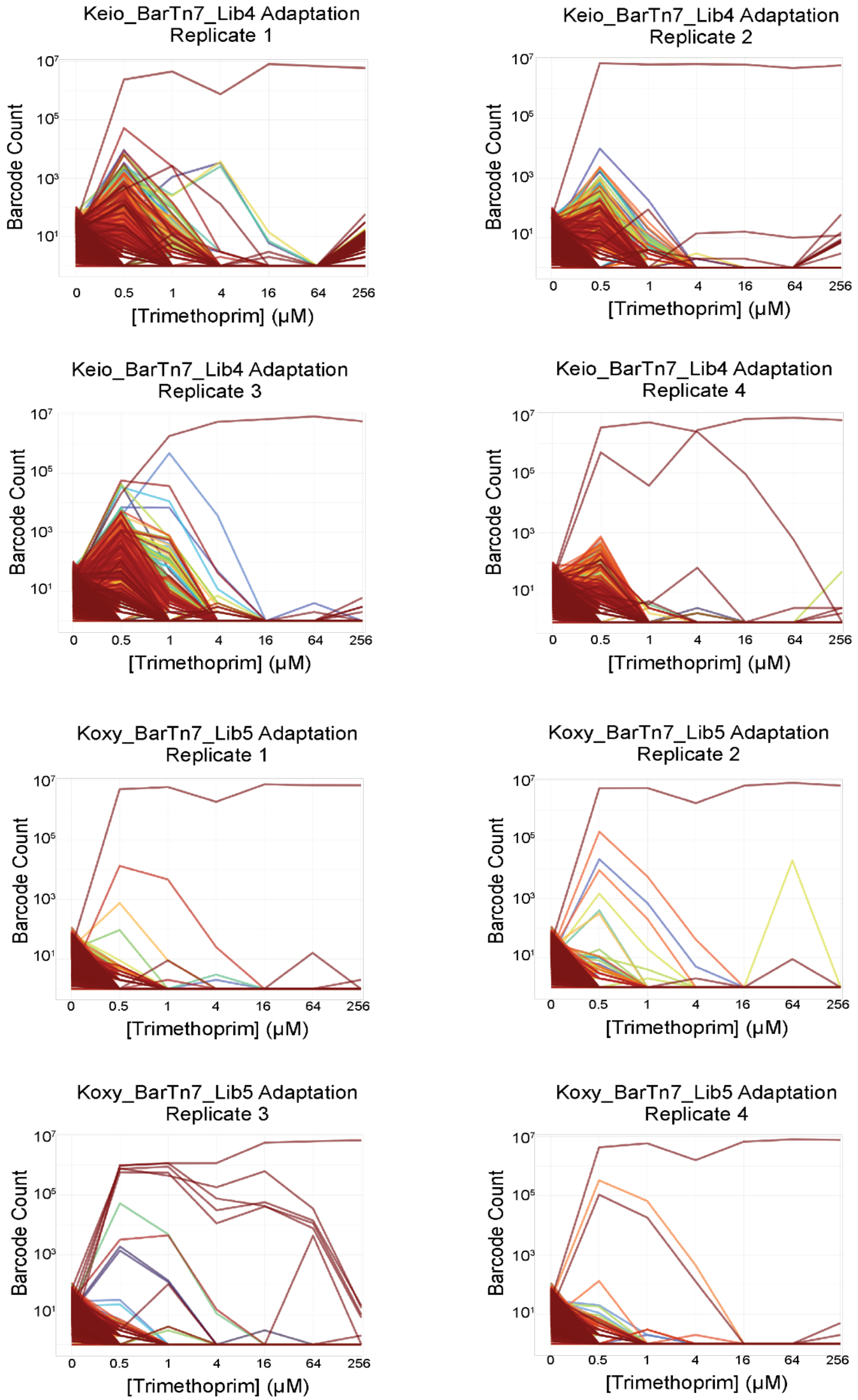

Keio_BarTn7_Lib4 Adaptation Replicate 1
Keio_BarTn7_Lib4 Adaptation Replicate 2
Keio_BarTn7_Lib4 Adaptation Replicate 3
Keio_BarTn7_Lib4 Adaptation Replicate 4
Koxy_BarTn7_Lib5 Adaptation Replicate 1
Koxy_BarTn7_Lib5 Adaptation Replicate 2
Koxy_BarTn7_Lib5 Adaptation Replicate 3
Koxy_BarTn7_Lib5 Adaptation Replicate 4
Barcode Count
$10^7$
$10^5$
$10^3$
$10^1$
0
0.5
1
4
16
64
256
[Trimethoprim] (μM)

**Figure S7.** Dynamics of low-abundance lineages during adaptation to trimethoprim. For each replicate adaptation of either Keio_BarTn7_Lib4 or Koxy_BarTn7_Lib5 to trimethoprim, each colored line represents a unique, barcoded lineage.
